# Single-cell RNA-sequencing reveals pre-meiotic X-chromosome dosage compensation in *Drosophila testis*

**DOI:** 10.1101/2021.02.05.429952

**Authors:** Evan Witt, Zhantao Shao, Chun Hu, Henry M. Krause, Li Zhao

## Abstract

Dosage compensation (DC) is a mechanism by which X chromosome transcription is equalized in the somatic cells of both males and females. In male fruit flies, expression levels of the X-chromosome are increased two-fold to compensate for their single X chromosome. In testis, dosage compensation is thought to cease during meiosis, however, the timing and degree of the resulting transcriptional suppression is difficult to separate from global meiotic downregulation of each chromosome. To address this, we analyzed testis single-cell RNA-sequencing (scRNA-seq) data from two *Drosophila melanogaster* strains. We found evidence that the X chromosome is equally transcriptionally active as autosomes in somatic and pre-meiotic cells, and less transcriptionally active than autosomes in meiotic and post-meiotic cells. In cells experiencing dosage compensation, close proximity to MSL (male-specific lethal) chromatin entry sites (CES) correlates with increased X chromosome transcription. We found low or undetectable level of germline expression of most *msl* genes, *mle, roX1* and *roX2* via sequencing or RNA-FISH, and no evidence of germline nuclear *roX1/2* localization. Our results suggest that, although DC occurs in somatic and premeiotic germ cells in *Drosophila* testis, there might be non-canonical factors involved in the dosage compensation. The single-cell expression patterns and enrichment statistics of detected genes can be explored interactively in our database: https://zhao.labapps.rockefeller.edu/gene-expr/.

## Introduction

In a wide variety of sexually reproducing animals, males and females have different numbers of X chromosomes. Somatic expression of X-linked genes need to be adjusted so that males and females produces similar levels of most proteins encoded on the variable chromosome (Prestel et al., 2010). Indeed, for many sexually reproducing animals, transcription is adjusted to compensate for differing numbers of sex chromosomes, a phenomenon called dosage compensation (Mukherjee and Beermann, 1965; Muller, 1950). Strategies for this process vary dramatically across the animal kingdom, some to increase chromosome X transcription in males, others to suppress it in females (Conrad and Akhtar, 2012; Kuroda et al., 2016; Mank et al., 2011). The spatial and temporal patterns of dosage compensation are key to understanding gene expression regulation and its roles in development. Single-cell sequencing has been successfully used to study dosage compensation during development (Mahadevaiah et al., 2020; Mahadevaraju et al., 2020), but has not yet been applied to adult spermatogenesis.

In somatic cells from male *Drosophila*, the DCC (including MSL-1, MSL-2 MSL-3, Mle, and Mof) acts in concert with Clamp (Soruco et al., 2013) and two noncoding RNAs, *roX1* and *roX2*, to increase transcription from the male X chromosome to levels comparable to the paired autosomes (Conrad and Akhtar, 2012; Franke and Baker, 1999; Samata and Akhtar, 2018). This binding is facilitated by the MSL proteins, which bind to a specific sequence motif in chromatin entry sites (CES) on the X chromosome and spread outward, upregulating local genes (Alekseyenko et al., 2008; Kelley et al., 1999). During male meiosis in some animal species, the sex chromosomes are downregulated in excess of that expected from the loss of dosage compensation. This is referred to as Meiotic Sex Chromosome Inactivation (MSCI) (Cloutier and Turner, 2010). Whereas the absence of dosage compensation would cause a 50 percent drop in X transcription, MSCI actively represses the X even further, leading to less than 50 percent X activity.

Dosage compensation in *Drosophila* somatic tissues (such as brain) has been extensively studied (Alekseyenko et al., 2008; Franke and Baker, 1999; Oh et al., 2003). However, whether dosage compensation occurs in germ cells (testis) and to what extent it plays a role in spermatogenesis, is actively debated. For instance, prior work has raised the possibility of germline dosage compensation (Gupta et al., 2006; Hense et al., 2007), although the magnitude and timing of this process are unclear. Other work suggests that demasculinization of the X chromosome might be partly due to dosage compensation in *Drosophila* (Bachtrog et al., 2010). However, Meiklejohn and Presgraves suggest that both dosage compensation and MSCI are absent during *Drosophila* spermatogenesis (Meiklejohn et al., 2011). Rastelli & Kuroda found that, unlike in somatic cells, the MLE protein does not associate specifically with the X chromosome in male germ cells, suggesting a possible alternative function of the *mle* gene in germ cells (Rastelli and Kuroda, 1998). Given that the MSL complex is thought not to localize to male germline X chromosomes (Rastelli and Kuroda, 1998), the mechanism of hypothetical germline DC is an additional mystery.

MSCI and dosage compensation are difficult to study in *Drosophila* because both somatic and sex chromosomes are downregulated during and after meiosis, making it challenging to identify groups of genes with deviant expression patterns in the context of global transcriptional downregulation. In some cases it may be difficult to distinguish the effect of X chromosome inactivation from the loss of dosage compensation, however, the multiple transgenic insertions in X and autosomes made by the Parsch group (Hense et al., 2007; Kemkemer et al., 2011) show that X inactivation exceeds that expected for loss of dosage compensation. Recent evidence supports spermatogonial dosage compensation in single-cell sequencing from larval testis (Mahadevaraju et al., 2020). We asked if the same is true in adult testis by analyzing gene expression from somatic, pre-meiotic, meiotic, and post-meiotic cells. We also asked if there was evidence of *roX1/2* activity in germline cells experiencing dosage compensation.

We sought to quantify the relative transcriptional dynamics of sex chromosomes and autosomes in a high throughput fashion with scRNA-seq of testis from two strains of *Drosophila melanogaster*. Compared to dissection-based methods, our methods allow high confidence in the identities of assigned germline cell types from somatic cell types, allowing us to holistically investigate germline X and autosome activity with greater precision. We found that pre-meiotic cells and somatic cells show X:autosome expression ratios close to 1, and GSC/early spermatogonia show evidence of excess dosage compensation (X:A = 1.83). After meiosis, X:autosome ratios decline to around 0.6-0.7, indicative of incomplete dosage compensation. Our observed pre-meiotic dosage compensation occurs despite negligible germline expression of *roX1* and *roX2*, which we confirmed with RNA-FISH. We also found that, in somatic and early germ cell types that have balanced X/autosome output, genes within 10,000 bp of an MSL CES have higher expression than genes further away. Our results support the existence of pre-meiotic X-chromosome dosage compensation in *Drosophila* testis via a noncanonical mechanism.

## Results

### Total transcription of X and autosomes peaks in early germ cells and is reduced after meiosis

We analyzed two single-cell testis RNA-seq datasets that we generated (Witt et al., 2019), totaling 13,000 cells. We chose to use two different strains to ensure that our findings are robust with respect to technical and strain variation, and repeated the analyses of each strain separately to ensure that all findings were reproducible (see methods). Using the ScTransform function from Seurat V3 (Satija et al., 2015), we combined data from two *D. melanogaster* strains and clustered corresponding cell types together on the same set of axes (Figure 1A, 1B). Consistent with our previous report, the two datasets correlate with a Pearson’s R of 0.97, and cells from both strains overlap well, with a qualitatively good distribution of cells from both strains around our dimensionally reduced dataset used for clustering (Witt et al., 2019) (Supplemental Figure 1). Compared to our previous report, in which we only used one strain of *D. melanogaster*, we were able to improve upon our previous cell type assignments by using a more accurate marker gene to denote cyst cells, *Rab11* (Joti et al., 2011) (Supplemental Figures 2, 3). Asking if the X and autosomes both follow expected trends of post-meiotic transcriptional downregulation, we counted the Unique Molecular Indices (UMIs, a proxy for RNA content) from X and autosomes in all cells. We observed that total RNA per cell from the X and autosomes peaks in late spermatogonia and early spermatocytes and is then reduced in late spermatogenesis (Figure 1C, 1D Supplemental Figure 4). If a cell type were dosage compensated, we would expect this ratio of X/A to be similar to or higher than that found in somatic cells. If a cell type lacked dosage compensation, we would expect this ratio to be lower than that of somatic cells. Taking a closer look, although the overall patterns for X and autosome look very similar (Figure 1C, 1D), the ratios of X/A are the highest in GSC and spermatogonia (0.120-0.145), intermediate in somatic hub, cyst and epithelial cells (0.104-0.135), and the lowest in meiotic (spermatocytes) and post-meiotic (spermatids) cells (0.087-0.090) (Supplemental Table 1). The relatively reduced ratio of X chromosome RNA in spermatocytes and spermatids suggests that the X chromosome is downregulated in excess of autosomes in meiotic and post-meiotic cells. The fact that this X/A ratio is higher in early germ cells than somatic cells indicates some form of excess dosage compensation.

**Table 1:** X and autosomal count ratios detected per gene for every cell type. These ratios are of raw counts from X and autosomes, not log transformed. The ratio of X/autosome expression is close to or greater than 1 in hub, cyst, GSC, early spermatogonia and late spermatogonia, indicating the presence of dosage compensation. This ratio decreases in meiotic and post-meiotic cells, indicating the loss of dosage compensation. P values are for a two-sided Wilcoxon test with a null hypothesis that X and autosome genes are similarly expressed, interpreted as dosage compensation. Meiotic and post-meiotic cells show an imbalance in x-autosome ratios, indicative of absent dosage compensation, but in pre-meiotic and somatic cells, x and autosomal ratios are near or over 1. The X/autosome ratio is highest in GSC/early spermatogonia, with a statistical enrichment of X chromosome gene expression, interpreted as possible excess dosage compensation in this cell type. DC means dosage compensation.

| Cell type | X/autosome ratio of raw counts | Raw p value | Adjusted P value | Interpretation |
| --- | --- | --- | --- | --- |
| Hub cells | 1.00 | 1.43e-01 | 2.86e-01 | DC |
| Cyst cells | 1.17 | 2.34e-02 | 8.24e-02 | DC |
| Epithelial cells | 0.92 | 1.63e-01 | 2.86e-01 | DC |
| GSC, Early spermatogonia | 1.83 | 6.44e-05 | 3.86e-04 | Excess DC |
| Late spermatogonia | 0.85 | 2.06e-02 | 8.24e-02 | DC |
| Early spermatocytes | 0.66 | 5.35e-08 | 4.82e-07 | No DC |
| Late spermatocytes | 0.63 | 6.69e-08 | 5.35e-07 | No DC |
| Early spermatids | 0.70 | 1.65e-05 | 1.16e-04 | No DC |
| Late spermatids | 0.77 | 1.30e-04 | 6.50e-04 | No DC |

**Figure 1:**
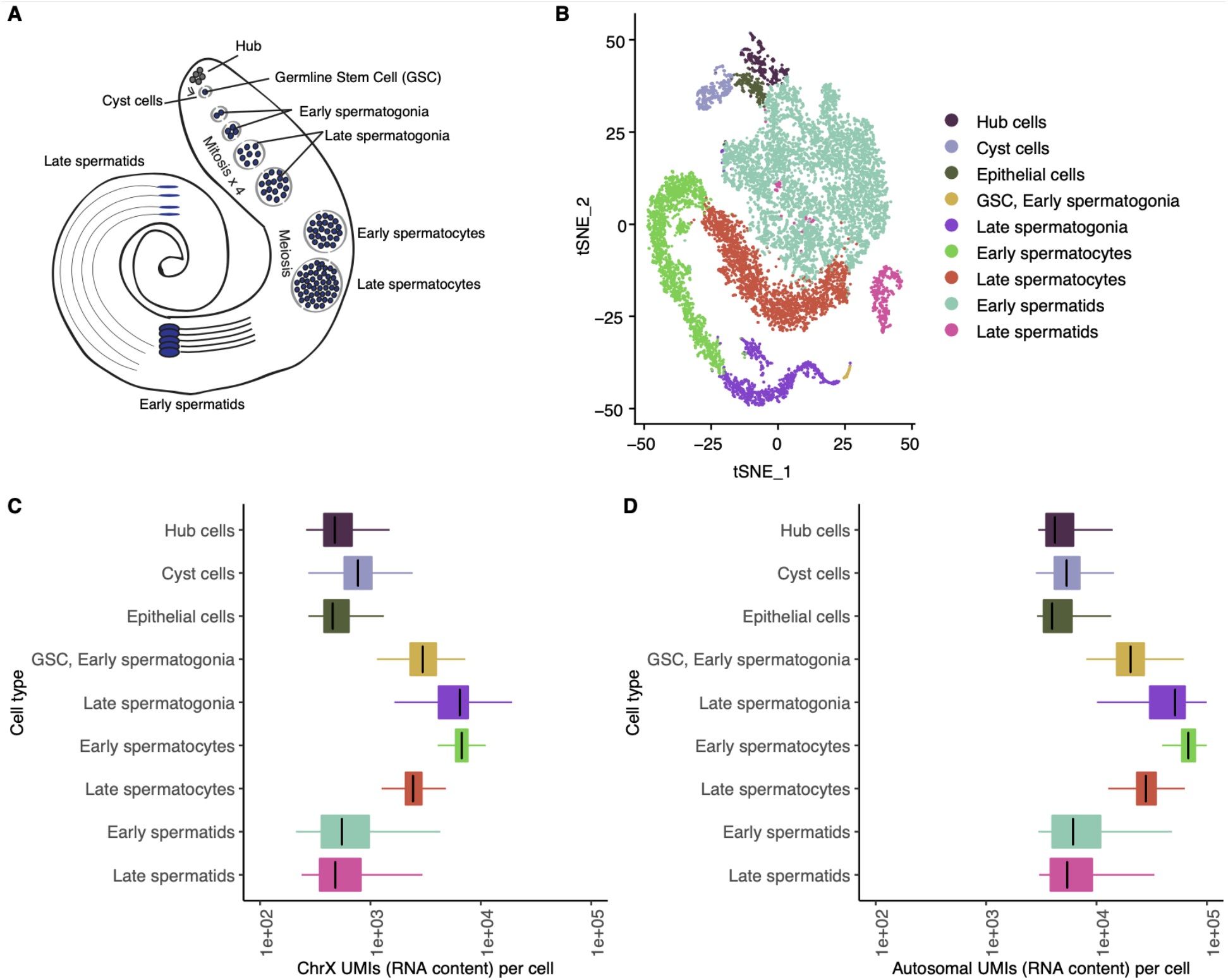
Integrated analysis of testis sequencing from two D. melanogaster strains. A.) A schematic showing the types of cells present inside *Drosophila* testis. Germline stem cells differentiate into spermatogonia, which undergo mitosis and become spermatocytes, which undergo meiosis and become haploid spermatids. B.) t-SNE plot showing a dimensional reduction of our combined, normalized libraries. Each point is a cell, clustered according to its similarity to other cells. C, D.) For every cell, the number of RNA molecules detected from the X chromosome and autosomes approximated by the summed number of Unique Molecular Indices counted per cell. X and autosomes are both downregulated during late spermatogenesis. Count numbers and ratios are shown in supplemental table 1.

**Figure 2:**
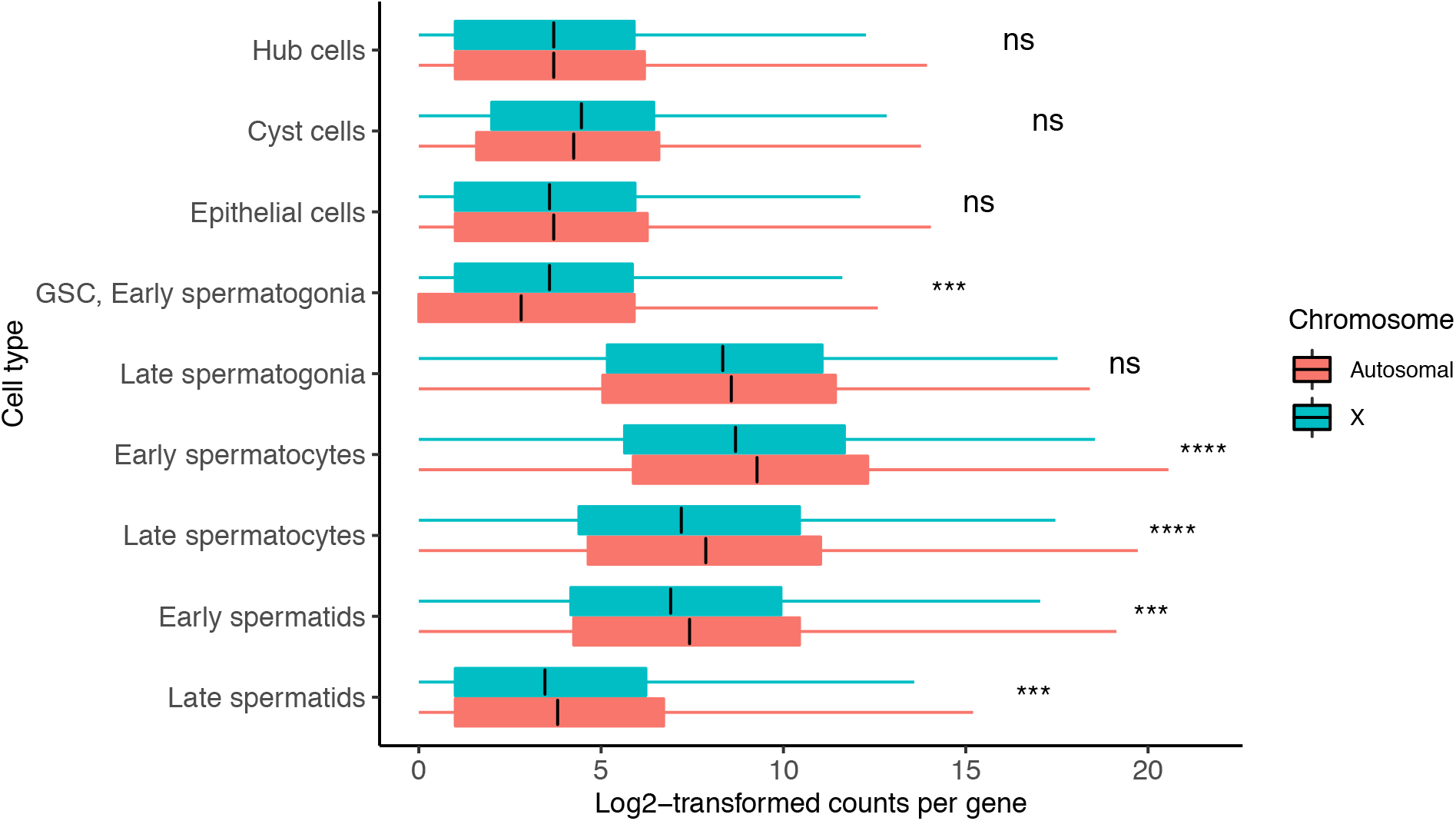
Dosage compensation equalizes X and autosomal transcription in select cell types. Shown is the median log-transformed (log2(counts+1) counts for every gene in X and autosomes. This ratio is similar in hub, cyst, epithelial cells, and late spermatogonia, indicating the presence of dosage compensation. GSC and early spermatogonia show a statistical enrichment of X chromosome transcription, indicating excess dosage compensation. Spermatocytes and spermatids have fewer X chromosomal than autosomal transcripts per gene, indicating that dosage compensation has been lost. This loss coincides with reduced expression of dosage compensation genes as seen in figure 2. Asterisks represent Holm-adjusted p values of a two-way Wilcoxon test of the null hypothesis that autosomal genes and X genes have equal detectable log-normalized counts in a cell type. In GSC, Early spermatogonia, we see a significant likelihood that the means of the two groups are not equal, and we can see that it is due to an enrichment of X chromosome counts. In meiotic cells this is the opposite-the significant value represents a depletion of X counts, since this is not a directional Wilcoxon test. Asterisks represent p values as follows: ns: >0.05, ^*^<0.05, ^**^<0.005, ^***^<0.0005, ^****^<0.00005. Detailed raw and adjusted p values, and interpretations of results of each cell type are in table 1.

**Figure 3:**
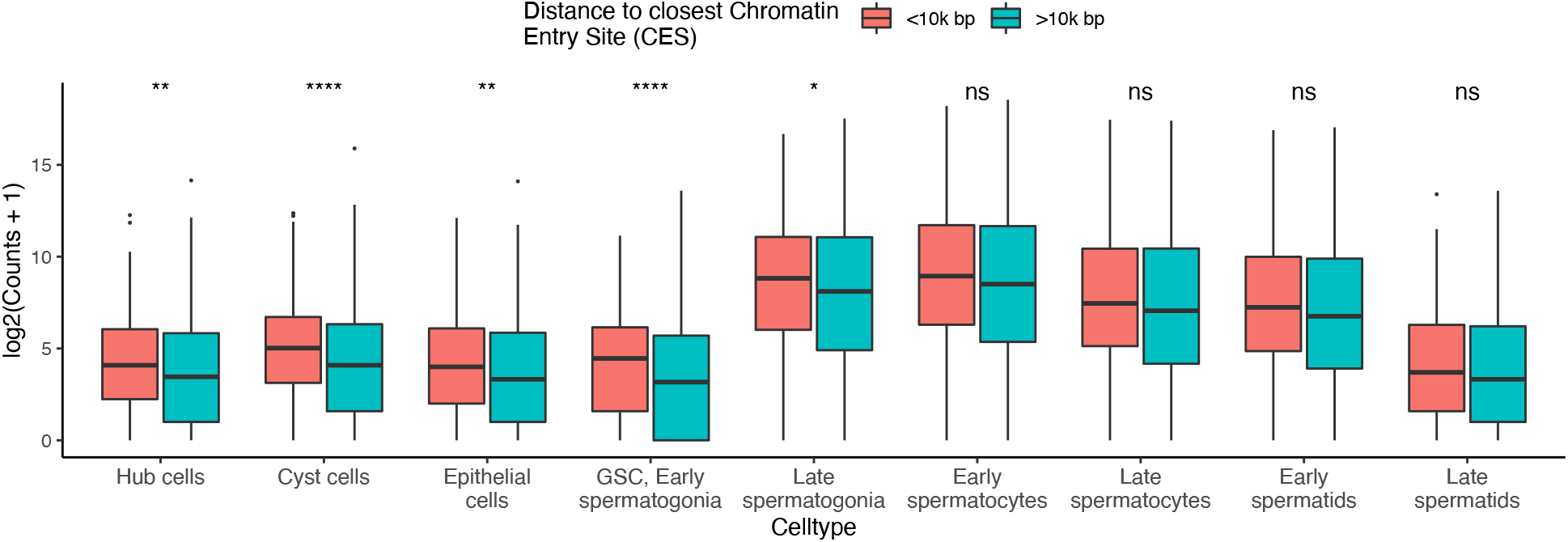
Close MSL CES correlates with increased transcription of X chromosome genes in cell types experiencing DC. The Y axis is log-transformed sum of counts for a given X chromosome gene in a cell type, grouped by proximity to a CES. Genes within 10,000 bp of a CES have statistically enriched expression compared to genes outside this range in all somatic cell types, and pre-meiotic germ cells (p values for two-tailed Wilcoxon test in table 2). These are the same cell types with statistically similar X and autosomal expression in figure 3. This indicates that the DCC is active in these cell types. Asterisks represent Holm-corrected p values as follows: ns: >0.05, ^*^<0.05, ^**^<0.005, ^***^<0.0005, ^****^<0.00005.

**Figure 4:**
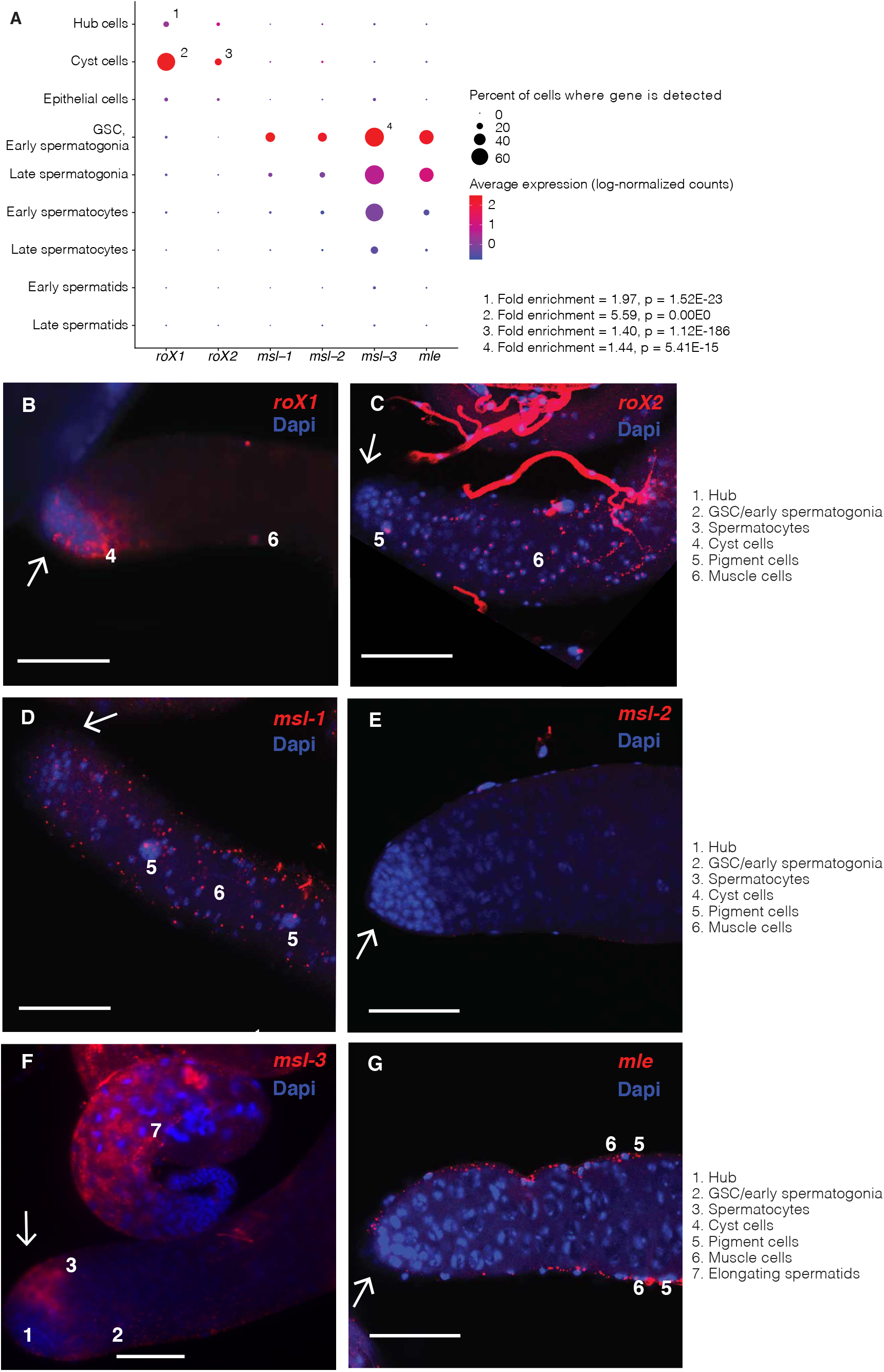
Expression patterns of DCC components in early germ cells. A. Expression patterns for DCC genes in testis scRNA-seq data. Dots are sized according to the percent of cells of a type where a transcript is detected, and colored according to expression (log-normalized counts) of that gene in the cell type. Numbers indicate that a gene is enriched in that cell type compared to all other cells. P values are adjusted with Bonferroni’s correction. *roX1* and *roX2* are enriched in cyst cells, *roX1* is enriched in hub cells, and *msl-3* is enriched in GSC/early spermatogonia. B-G) RNA-FISH of the distal testis region. White arrows indicate the hub region. Scale bars are 50 µM for all images. B) *roX1* is usually but not always found in somatic cyst cells near the testes tip. Later, it is expressed in elongating spermatid cysts, and in the nuclei of the somatic terminal epithelial cells (not shown). No nuclear expression was detected in germline cells at the distal tip (arrow). ScRNA-seq indicates expression in somatic cyst cells, but no significant germline expression. C) in FISH, *roX2* shows nuclear localization in peripheral muscle and pigment cells, with cytoplasmic expression detected in elongating spermatid cysts. Little to no expression was observed in the germ cell region. In scRNA-seq it shows 0 detectable counts in most cells of every type, but detectable counts more often in somatic cells than germ cells. D) in FISH, *msl-1 also* shows some expression in muscle and pigment cells at the testis periphery, but no such foci inside the testis. In scRNA-seq, by contrast, transcripts are faintly detectable in GSC/early spermatogonia. E) in FISH, *msl-2* was undetectable, but at higher exposure showed faint cytoplasmic expression throughout the region, and scRNA-seq finds weak, stochastic expression in early germ cells. F) in FISH, *msl-3* shows weak expression and nuclear localization in hub cells, early germ cells and elongating spermatids, as well as in the nuclei of muscle and pigment cells. In scRNA-seq data it is stochastically expressed in somatic cells but commonly expressed in early germ cells. Nuclear localization in hub, muscle, pigment and early germ cells is shown in Supplemental Figure 14. G) in FISH *mle* shows bright foci in muscle and pigment cells but only weak expression in early stem cells (image is focused on peripheral cells at the testes edges and testis lumen). In scRNA-seq, expression levels peak in GSC/early spermatogonia and decline thereafter.

### Relative RNA production from the X chromosome indicates dosage compensation in pre-meiotic cells

Since the total RNA content of a cell is an imperfect measure of gene expression trends, we asked whether the RNA content produced individual genes from the X chromosome and autosomes is roughly equivalent in different stages of spermatogenesis. To ask whether a stage favors X or autosomes, we counted the detected reads per gene in each cell type for X and autosomes. If a cell type produces roughly equal amounts of RNA from X and autosomal genes, we would consider that cell type subject to dosage compensation. If a cell type produced less RNA from a median X gene than an autosomal gene, we would interpret this as the absence of dosage compensation, similar to earlier work (Kemkemer et al., 2011).

**Table 2:**
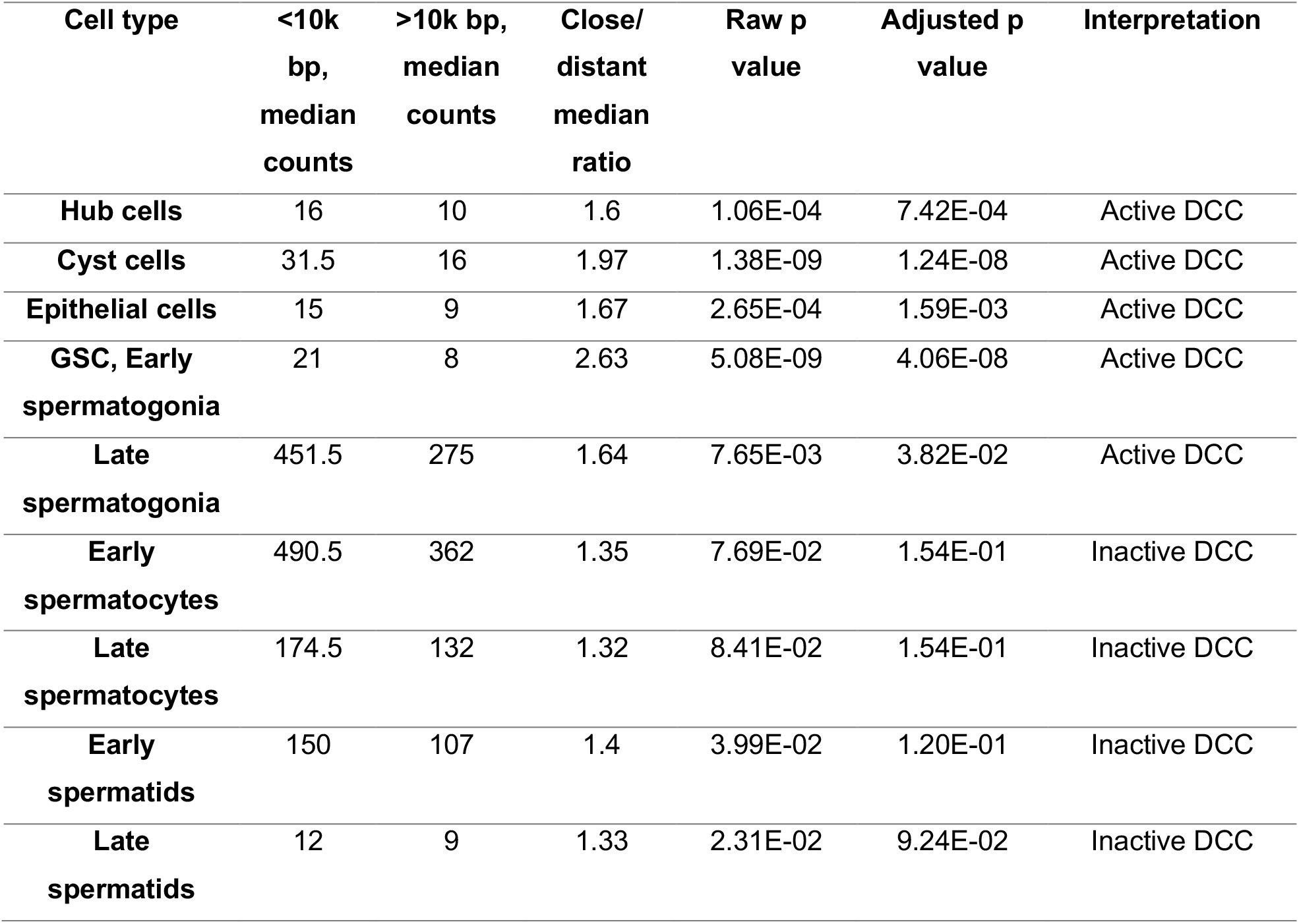
Evidence of DCC activity in somatic cells and pre-meiotic germ cells. In each cell type we have calculated the median counts for X chromosome genes within 10000 bp of an annotated CES and genes outside 10000 bp, corresponding to Figure 4. In hub, cyst, epithelial, GSC, early and late spermatogonial cells, a two-tailed Wilcoxon test finds that these two sets of genes have statistically different count distributions with an alpha of 0.05. These cell types have a higher ratio of median counts from close and distant genes, indicating that CES sites influence X chromosome gene expression in these cell types, a hallmark of dosage compensation. The cell type with the highest difference between close/distant x gene expression is GSC/early spermatogonia, which also had the highest X/autosome ratio in table 1.

| Cell type | <10k bp, median counts | >10k bp, median counts | Close/ distant median ratio | Raw p value | Adjusted p value | Interpretation |
| --- | --- | --- | --- | --- | --- | --- |
| Hub cells | 16 | 10 | 1.6 | 1.06E-04 | 7.42E-04 | Active DCC |
| Cyst cells | 31.5 | 16 | 1.97 | 1.38E-09 | 1.24E-08 | Active DCC |
| Epithelial cells | 15 | 9 | 1.67 | 2.65E-04 | 1.59E-03 | Active DCC |
| GSC, Early spermatogonia | 21 | 8 | 2.63 | 5.08E-09 | 4.06E-08 | Active DCC |
| Late spermatogonia | 451.5 | 275 | 1.64 | 7.65E-03 | 3.82E-02 | Active DCC |
| Early spermatocytes | 490.5 | 362 | 1.35 | 7.69E-02 | 1.54E-01 | Inactive DCC |
| Late spermatocytes | 174.5 | 132 | 1.32 | 8.41E-02 | 1.54E-01 | Inactive DCC |
| Early spermatids | 150 | 107 | 1.4 | 3.99E-02 | 1.20E-01 | Inactive DCC |
| Late spermatids | 12 | 9 | 1.33 | 2.31E-02 | 9.24E-02 | Inactive DCC |

In meiotic and post-meiotic cells, we observe a depletion in the relative RNA produced from the X chromosome. We found that the median reads per gene were lower for the X than autosomes in early and late spermatocytes, and early and late spermatids in both fly strains (Figure 2, Table 1, Supplemental Figure 5, Holm-adjusted p values: 4.82e-07, 5.35e-07,1.16e-04 and 6.50e-04, respectively). Interestingly, X and autosomes produced roughly equal numbers of reads per gene in somatic hub, cyst, and epithelial cells, and germline late spermatogonia (adjusted p values: 2.86e-01, 8.24e-02, 2.86e-01, 8.24e-02, respectively). In GSC and early spermatogonia, the X:autosome ratio increases to 1.83, suggestive of excess dosage compensation (adjusted p value: 3.86e-04). We repeated this analysis with gene expression values linearly scaled from 0 to 1 and found similar results (Supplemental Figure 6).

### Gene expression is enriched for genes close to an MSL CES

The DCC is thought to facilitate dosage compensation by binding to specific DNA motifs and spreading outward, upregulating local genes (Alekseyenko et al., 2008). We examined transcriptional activity for every gene approximated by the total counts detected in a cell type and found evidence that, in certain cell types, genes close to a CES have higher RNA output (counts) than genes further away (Figure 3, Supplemental Figure 7,8, Table 2, Supplemental Table 2). These cell types are hub, cyst, epithelial, GSC, early spermatogonia and late spermatogonia (Adjusted p values for two-tailed Wilcoxon test=7.42e-04, 1.24e-08, 1.59e-03, 4.06e-08, and 3.82e-02, respectively) the same cell types for which we observed evidence of dosage compensation in Figure 2. In cell types without evidence of DC, there is not a significant difference in detectable counts per gene between genes <10000 bp from a CES, and genes further away. The same trend is present in somatic cells for which we observed dosage compensation, and also in pre-meiotic germ cells. Pearson’s correlations between distance and expressed counts are higher for somatic and pre-meiotic cells than meiotic and post meiotic cells (Supplemental Figure 9). This suggests that our observed dosage compensation in somatic and pre-meiotic cells is mediated from the same sequence element, with possible different mechanisms.

### Testis-biased and testis-specific genes do not bias apparent dosage compensation

To understand whether testis biased genes influence the observed patterns of dosage compensation, we repeated the analyses from Figures 1,2 and 3 after removing testis-specific or testis-biased genes from our data. Supplemental Figures 8, 10, and 11 show that the appearance of dosage compensation in somatic and pre-meiotic cells remains without these genes. One difference is that when testis-specific genes are removed from the analysis, there is no longer evidence of excess dosage compensation in GSC/early spermatogonia (Supplemental Figure 11), indicating that these genes are disproportionately upregulated in these cells. Without these testis-specific genes, the distributions of X and autosome counts are similar, consistent with the degree of dosage compensation observed in somatic cells. This could indicate an uneven degree of dosage compensation for different gene classes in early germ cells. Whether the excessive up-regulation of testis-biased genes is through dosage compensation or other mechanism is unclear, however, it would be important to study the mechanism and the impact of this pattern in the future.

### Germline dosage compensation takes place despite low levels of *roX1* and *roX2*

We sought to explain our observed testis somatic and premeiotic dosage compensation by querying the expression patterns of genes known to regulate this process in somatic cells. The DCC contains proteins such as *msl1, msl2, msl-3, mof, mle*, and *Clamp*, and two ncRNA’s, *roX1* and *roX2*. We examined the Flyatlas2 fly tissue expression data (Leader et al., 2018), and found that the ncRNAs *roX1* and *roX2* are depleted several fold in testis compared to most other tissues, while transcripts of the protein components of this complex are expressed more uniformly (Supplemental Figure 12).

In our single-cell data, we found low-level expression of transcripts from protein components of the DCC in early germ cells but found them less expressed in somatic cells (Figure 4A). The only germline enrichments of any DCC components were *Clamp* and *msl-3* in GSC/early spermatogonia (adjusted p values 1.38E-09 and 5.41E-15, respectively). Conversely, we detected robust *roX1* enrichment in somatic hub and cyst cells (adjusted p values 1.52E-23, 0.00E00, respectively), and *rox2* enrichment in cyst cells (adjusted p value 1.12E-186), but germline expression was undetectable, concordant with the *roX* gene depletion observed by Flyatlas2 (Supplemental Figure 12).

To confirm these findings, we performed RNA fluorescent in situ hybridization (RNA-FISH) on *roX1, roX2, msl-1, msl-2, msl-3*, and *mle*. RNA-seq and scRNA-seq data suggest that the expression levels of the DCC genes differ at the tissue and cell-type level, consistent with the RNA-FISH results. For example, *msl-3* is the most highly expressed component of the DCC in RNA-FISH, whole-tissue RNA-seq, and scRNA-seq. *Mle* is detectable in the early germline in scRNA-seq, but we could not detect early germline *mle* expression with FISH. In FISH, we saw *mle* expression in peripheral muscle and pigment cells, concordant with its canonical role in somatic dosage compensation. The whole-tissue RNA depletion of *roX1* and *roX2* coincides with low/sparse expression in our testis scRNA-seq data as well as FISH images. FISH did not reveal significant germline expression of any *msl* gene except msl-3. Many DCC genes such as *roX2, msl-1, msl-2*, and *msl-3* showed distinct nuclear localization in somatic muscle and pigment cells, which were not captured by scRNA-seq.

To verify that the apparent absence of DCC transcripts in the germline was biological, not technical, we also performed RNA-FISH on *Drosophila* accessory glands with the same probes. All of the *msl* genes, as well as *roX1, roX2*, and *mle*, show distinct foci and nuclear RNA localization in accessory gland RNA-FISH (Supplemental Figure 13).

## Discussion

In this work, we directly observe dosage compensation in terms of X:autosome count ratios and differential gene activity around CE sites. These lines of evidence point to adult pre-meiotic dosage compensation for the X chromosome, consistent with recent findings in larval testis (Mahadevaraju et al., 2020). After meiosis, the X chromosome is downregulated to an extent that would be expected from incomplete dosage compensation. Compared to prior methods which use transgenic reporters to query expression from various autosomal and sex chromosome loci (Hense et al., 2007; Kemkemer et al., 2011), our approach provides a direct, holistic and high-throughput picture of whole-chromosome transcriptional trends in each cell type.

We found evidence for dosage compensation in three types of somatic cells and two types of pre-meiotic germ cells. These cells, however, express a different repertoire of DCC genes. Some somatic cells express *roX1* or *roX2*, but show sparse quantities of the *msl* genes and *mle*, other mediators of somatic dosage compensation. In contrast, premeiotic germ cells showed some expression of *msl-3* but *roX1* and *roX2* are barely detectable. This is concordant with depletion of *roX, msl-1*, and *msl-2* RNA in testis compared to other tissues in Flyatlas2 RNA-seq data. However, RNA levels of the DCC genes do not correspond to the actual magnitude of dosage compensation in the adult male germline, suggesting that abundances of proteins and RNAs in the cells may not be well-correlated. Enriched gene expression around MSL CES sites in these cell types strongly implies some sort of active DCC in somatic and early germ cells. This would normally indicate the activity of the DCC, but we were unable to detect significant germline expression of several components of the DCC with scRNA-seq or RNA-FISH. While the DCC proteins could be present and active without active transcription, the lack of germline *roX* expression is a more reliable clue for an alternative mechanism, since these are ncRNAs. In our scRNA-seq data, no DCC genes were enriched in germ cells except *msl-3* and *Clamp*, which were enriched in GSC/early spermatogonia (Figure 4A). Rather than acting in dosage compensation, *msl-3* might instead be facilitating entry into meiosis, as suggested by recent work in mice (McCarthy et al., 2019). If so, this could suggest a conserved function of *msl-3* across kingdoms and explain the lack of germline enrichment of any other DCC genes in our data.

While it is possible that a cell could perform DC without active transcription of DCC proteins, we would expect to find evidence of germline nuclear localization of *roX1* and *roX2* ncRNAs as we see in accessory glands. With RNA-FISH and scRNA-seq, we found evidence of germline *msl-3* expression, however, most of the other DCC components were so lowly expressed in the germline that we could only detect a few counts per cell with sequencing, and no enrichment with RNA-FISH. Our results do not rule out possible low levels of DCC proteins being present in pre-meiotic cells, but strongly demonstrate the absence of *roX1* and *roX2* in these cells. RNA-FISH suggests that the outer somatic layers of the testis express most DCC genes, so their apparent depletion in scRNA-seq may be due to underrepresentation of the somatic pigment and muscle cells of the testis sheath.

Our results suggests that germline dosage compensation might occur without all of the canonical components of the DCC, consistent with earlier studies (Gupta et al., 2006; Meller and Rattner, 2002; Rastelli and Kuroda, 1998). There could also be germline-specific unannotated isoforms of the *msl* genes that evade detection with RNA probes, gene annotations or antibodies tailored for variants discovered in somatic cells. Alternatively, the permissive chromatin environment of the testis (Soumillon et al., 2013) may be more conducive to dosage compensation, requiring only small amounts of *roX1* and *roX2* to boost X transcription to autosomal levels, levels that evade detection with RNA-FISH. On the other hand, Rastelli and Kuroda (Rastelli and Kuroda, 1998) reported that H4K16ac is not enriched on the X in testes germ cells and the MSL proteins are undetectable in germ cells with immunostaining. This could indicate that the dosage compensation we observe may occur independently of the canonical DCC.

Since we observed likely germline activity of MSL CE sites, it is possible that there exists a noncanonical mechanism of early germline DC that uses the same sequence elements as somatic cells. It may perhaps be mediated by CLAMP, which has been shown to bind to MSL CE sites and increase X chromosome accessibility (Urban et al., 2017), and which we observed upregulated in early germ cells (Figure 4). There is also a possibility that MSL recruitment is associated with other factors such as DNA replication (Lucchesi and Kuroda, 2015), thus the MSL recruitment and the timing of dosage compensation in cells can be very dynamic.

Our results do not provide evidence to support or disprove MSCI in late germ cells. We observed, in spermatocytes and spermatids, that X chromosome genes are expressed with median RNA counts of 63-77 percent that of autosomal genes. This would be expected from incomplete or absent dosage compensation, as some transcripts present in these cells would be retained from earlier stages despite reduced X transcription. For us to confidently predict the presence of MSCI, we would have to observe X:autosome ratios <0.5 in meiotic and postmeiotic germ cells. While X:autosome ratios rise from 0.63 in late spermatocytes to 0.77 in late spermatids, this could be driven by reduced transcription of both the X and autosomes making leftover transcripts proportionally more abundant.

While many exciting mechanistic questions remain, our study strongly supports the presence of dosage compensation in *Drosophila* premeiotic germ cells, and shows that it occurs despite sparse germline expression of *roX1, roX2* and most DCC proteins. Pre-meiotic dosage compensation appears to be driven from the same chromatin elements as somatic dosage compensation. Future studies should confirm that our observed trends of dosage compensation in scRNA-seq are abrogated when MSL CE sites are blocked, or that they occur during a testis-specific knockdown of *roX1* and *roX2*.

## Methods

### Integration and normalization of *D. melanogaster* scRNA-seq datasets

Raw reads from the two datasets were aligned to the FlyBase dmel_r6.15 reference GTF and Fasta (Thurmond et al., 2019) with Cellranger. The count matrices were made into separate Seurat objects, normalized and scaled with default Seurat parameters, and integrated with Seurat SCtransform. Neighbor finding, clustering and dimensional reduction were performed on the combined dataset. The exact parameters and commands used are available in the accompanying GitHub repository (see Data Availability). For all analyses, genes are only considered if they are detected in at least 3 cells, and cells are only considered if they express >200 genes. A gene is considered “expressed” in a cell if it has at least 1 read in that cell. Expression for figure 4 was calculated with the NormalizeData function of Seurat, with a scale factor of 10000.

### Assigning cell types from the integrated single-cell dataset

Cell types were assigned mostly the same marker genes from Witt et. al 2019. Clusters expressing *bam* and *aub* were assigned as a mixture of germline stem cells (Rojas-Ríos et al., 2017) and early spermatogonia (Kawase, 2004). Clusters with slight enrichment of *bam*, but less *His2Av* than GSC/early spermatogonia were interpreted as late spermatogonia. This was corroborated by the fact that this group is adjacent to the GSC/early spermatogonia cluster in the t-SNE. As we previously found that *fzo* expression peaks earlier than *twe* in spermatocytes, we assigned clusters with the highest enrichment of *fzo* (Hwa et al., 2002) as early spermatocytes, and cells with *twe* (Courtot et al., 1992) but no *fzo* as late spermatocytes (Witt et al., 2019). We defined early spermatids as clusters with enriched *Dpy-30L2* but no *fzo* or *twe*, and clusters enriched for *p-cup* as late spermatids (Barreau et al., 2008). Cells enriched for *Fas3* were deemed somatic hub cells, clusters enriched for *zfh-1* were assigned as somatic cyst cells (Zhao et al., 2009), and cells enriched for *MtnA* (Faisal et al., 2014) but not *Fas3* were labelled somatic epithelial cells. One major update from our 2019 paper is that we used *Rab11*(Joti et al., 2011), not zfh1, as a marker gene for cyst cells, improving the cell-type assignments.

### Calculating relative RNA content from each cell type

RNA content per cell is the sum of all Unique Molecular Indices (UMIs/counts) detected from the X chromosome or autosomes in a cell type, divided by the number of cells. This is a proxy for the relative RNA content per cell and is not a measurement of the actual number of RNA molecules present.

### Comparing RNA output by chromosome and spermatogenic stage

For this method, we obtained the total RNA counts from every X and autosomal gene in every cell type. We then log transformed these counts with y= Log2(counts+1) and performed non-directional Wilcoxon tests, with Holm-corrected p values indicating if genes from the X chromosome are likely to have equal median counts to genes from the autosomes.

### Correlating chromatin entry sites with nearby gene output

We obtained a list of MSL recognition sites (Alekseyenko et al., 2008) and converted the coordinates to *D. melanogaster* version 6 with the FlyBase coordinates converter (Thurmond et al., 2019)(Supplemental Table 3). For every X chromosome gene, we then calculated the distance between the start coordinate of its gene region and the start coordinate of the closest chromatin entry sites. For each gene, we summed all the reads detected in each cell type. We log transformed these counts for every cell type with the equation Log2(counts+1) and calculated Pearson’s R and corresponding p values between distance and gene counts for every cell type. We then adjusted these p values with Holm’s method.

### RNA-FISH of DCC transcripts in *Drosophila* testis

Plasmids from Drosophila Gene Collection (DGC) libraries were used to generate Rox1 (CR32777), Mle (CG11680), MSL-1 (CG10385), MSL-2 (GH22488), MSL-3 (CG8631) DIG-labeled probes. The following primers were designed for Rox2 (CR32665) template production by PCR of genomic DNA: forward: 5’-AATTAACCCTCACTAAAGGGT T GCCATCGAAAGGGTAAATTG-3’, reverse: 5’-GTA ATA CGA CTC ACT ATA GGG CAGTTTGCATTGCGACTTGT-3’. RNA probe preparation and use were carried out as previously described (Wilk et al, 2017; Jandura et al., 2017). Images were acquired using a Leica Sp8 confocal microscope.

### Calculating transcript enrichment in scRNA-seq data

For the dosage compensation genes, we used the FindAllMarkers Seurat function to identify genes enriched in a cell type compared to all other cells. We adjusted P values with Bonferroni’s correction.

## Abbreviations and acronyms

scRNA-seq: single-cell RNA-sequencing
RNA-seq: RNA-sequencing
MSL: male-specific lethal
DC: dosage compensation
DCC: dosage compensation complex
CES: chromatin entry sites
MSCI: Meiotic Sex Chromosome Inactivation
RNA-FISH: RNA fluorescent in situ hybridization

## Reproducibility

To ensure that our findings are robust with respect to biological and technical variability, we performed analyses from the main manuscript separately on each strain. The main findings from the paper are highly similar in both strains, with only small differences that don’t negate our main findings. These results are in Supplemental Figures 3, 4, 5, and 7.

## Data availability

R Code for integrating the Seurat objects for the 517 and wild strains is available at https://github.com/LiZhaoLab/Singlecell_DosageCompensation. Code has also been deposited for the mathematical analyses and generation of figures. Fastq files of the single-cell testis RNA-seq data have been deposited at NCBI SRA with accession numbers SAMN10840721 (RAL517 strain, BioProject # PRJNA517685) and SAMN12046583 (Wild strain, PRJNA548742). The integrated Seurat object containing raw counts, calculated expression, cell type assignments and strain information is available at https://rockefeller.box.com/s/7t5s060t7olewlpwmtjlef2ifc45u1wu, along with data files needed to complete the analysis in the R markdown file on GitHub. The expression pattern and statistics of each gene can be found in the database: https://zhao.labapps.rockefeller.edu/gene-expr/.

## Acknowledgment

We thank Zhao lab members for useful discussion and critical reading of the manuscript, especially Nicolas Svetec for providing essential ideas. The authors would like to thank the reviewers whose comments improved the quality of the work.

