## supplemental figures and tables for "Single-cell RNA-sequencing reveals pre-meiotic X-chromosome dosage compensation in *Drosophila testis*"

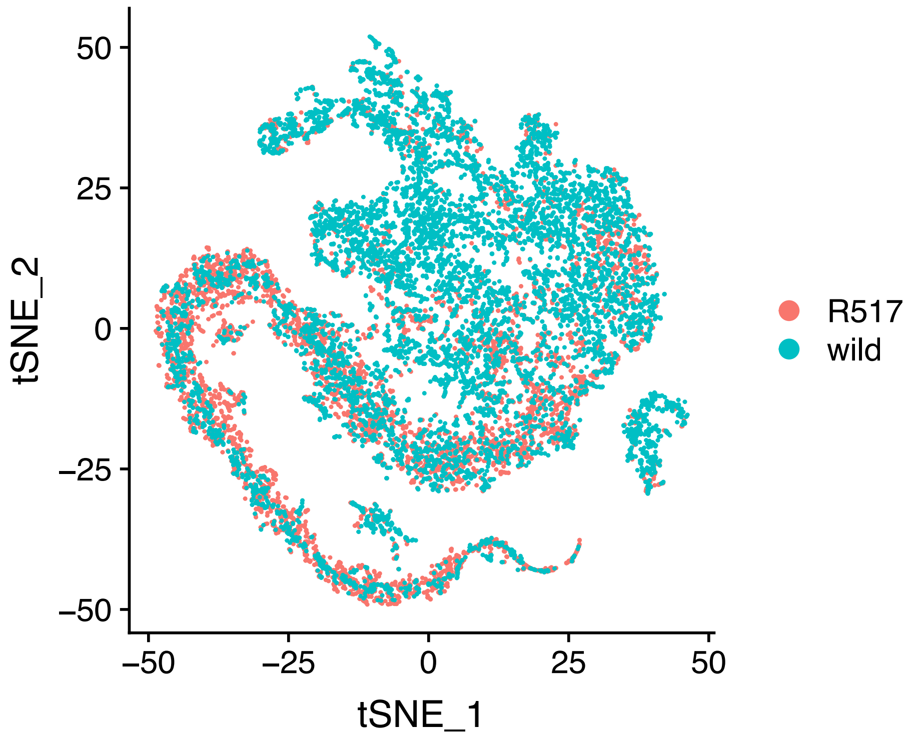


**Supplemental Figure 1:** Integrated t-SNE of both strains corresponding to figure 1A. Both strains overlap fairly well. Cell types were assigned from this integrated dataset.

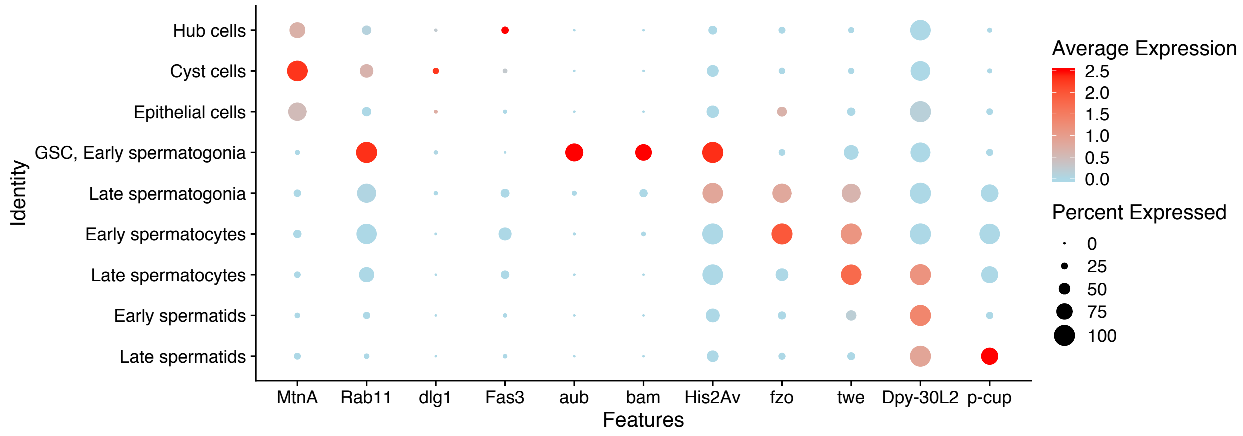


**Supplemental Figure 2: Relative enrichment of marker genes used to assign cell types.** These genes were used to assign cell types, with details in the methods section.

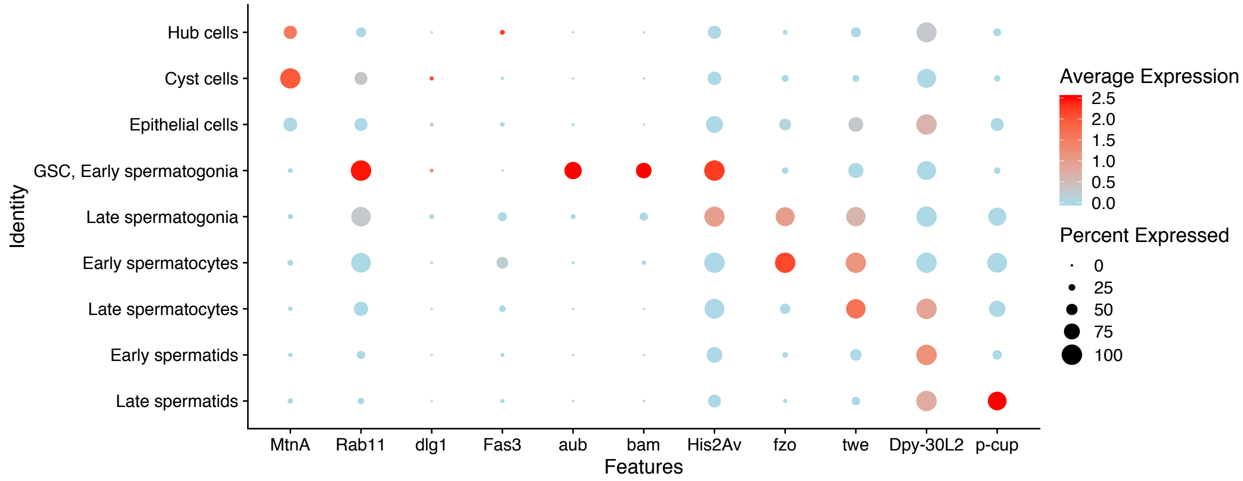


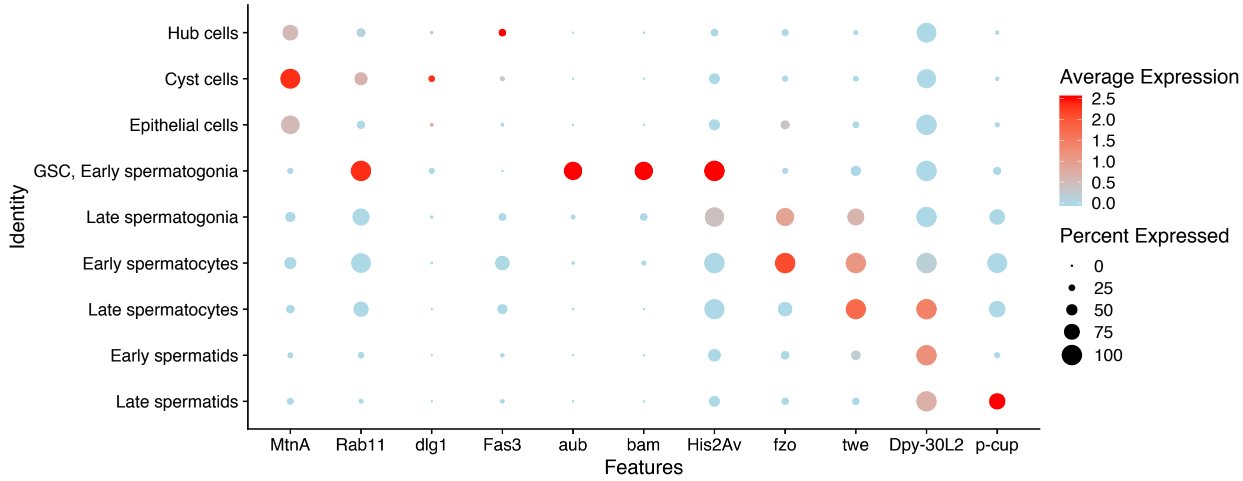


**Supplemental Figure 3: Marker genes used to assign cell types, split by strain**. This corresponds to supplemental figure 2. Marker enrichment in both strains corroborates cell type assignments.

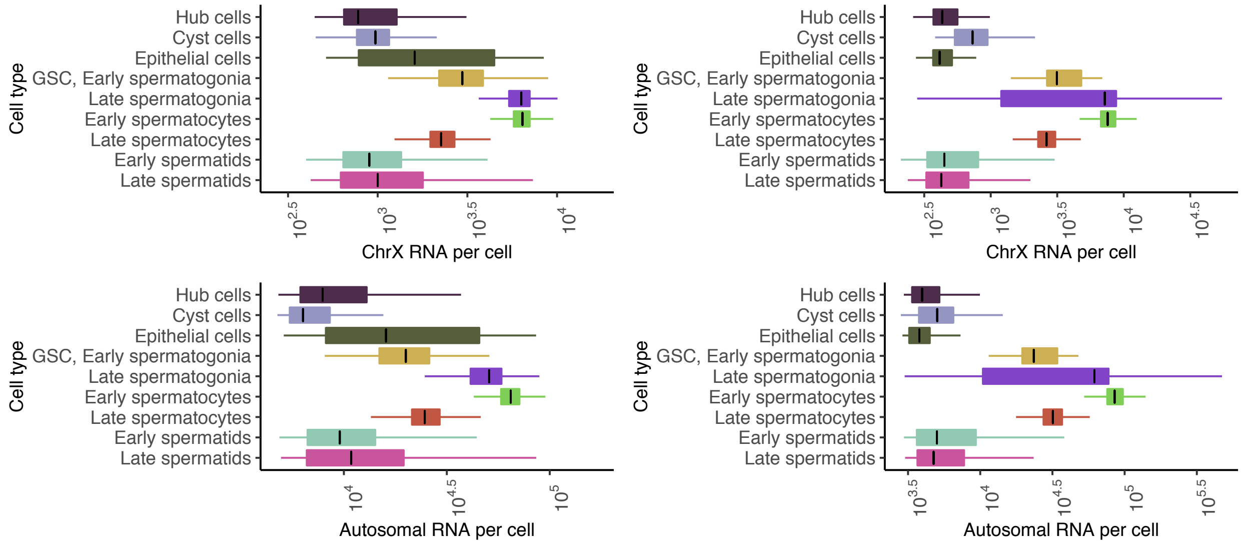


**Supplemental Figure 4:** RNA per cell from X and autosomes, for each of the two strains separately (Left is R517, right is wild). This corresponds to Figure 1C and 1D.

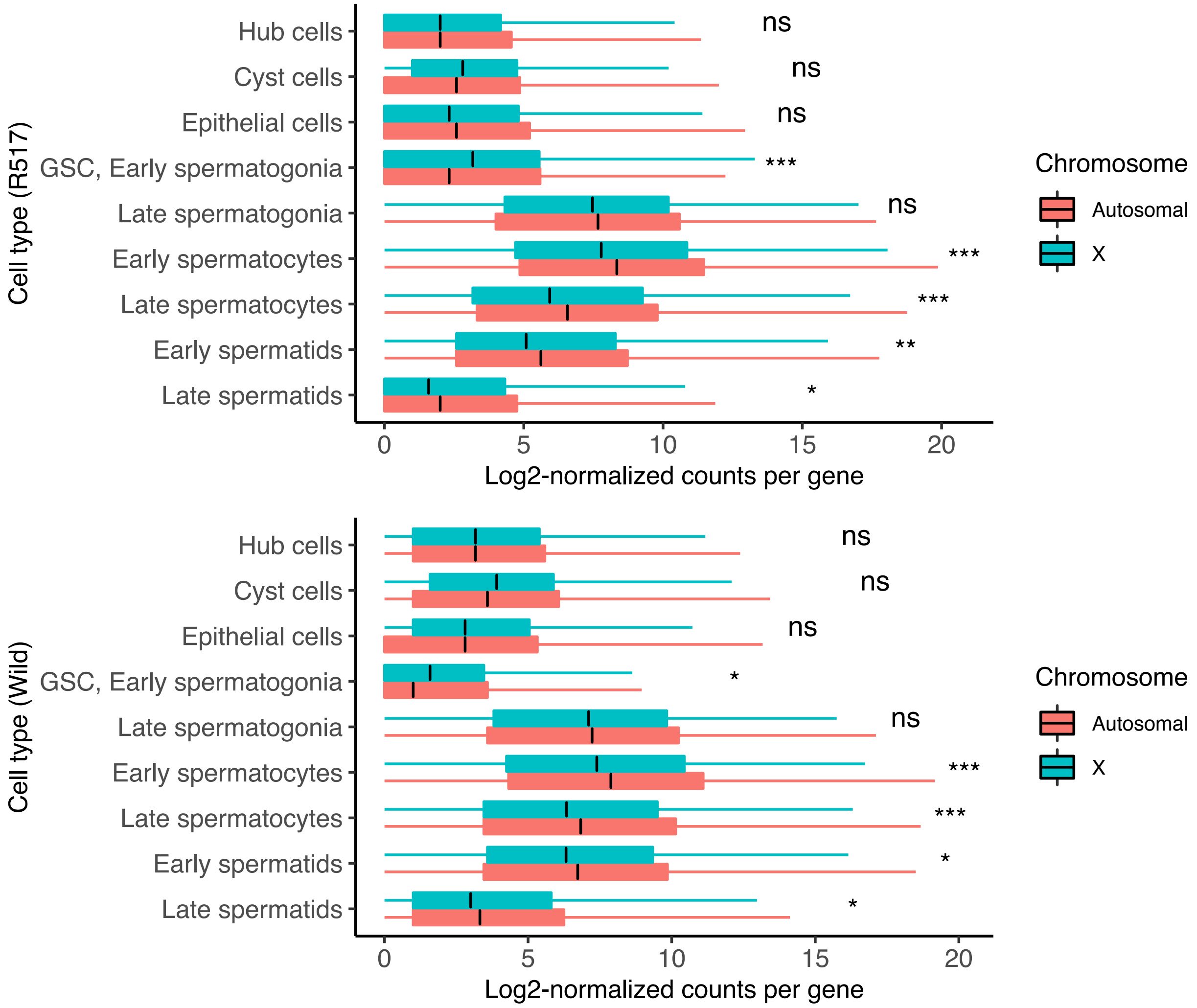


**Supplemental Figure 5: Median counts from X and autosome, split by strain.** Corresponding to figure 2, both strains support the appearance of pre-meiotic dosage compensation, somatic dosage compensation, and meiotic and post-meiotic X downregulation. In addition, in both strains, GSC and early spermatogonia appear to show X over-compensation. P values are from a two-tailed Wilcoxon test of the null hypothesis that X and autosomal counts are equal, adjusted with Holm's correction.

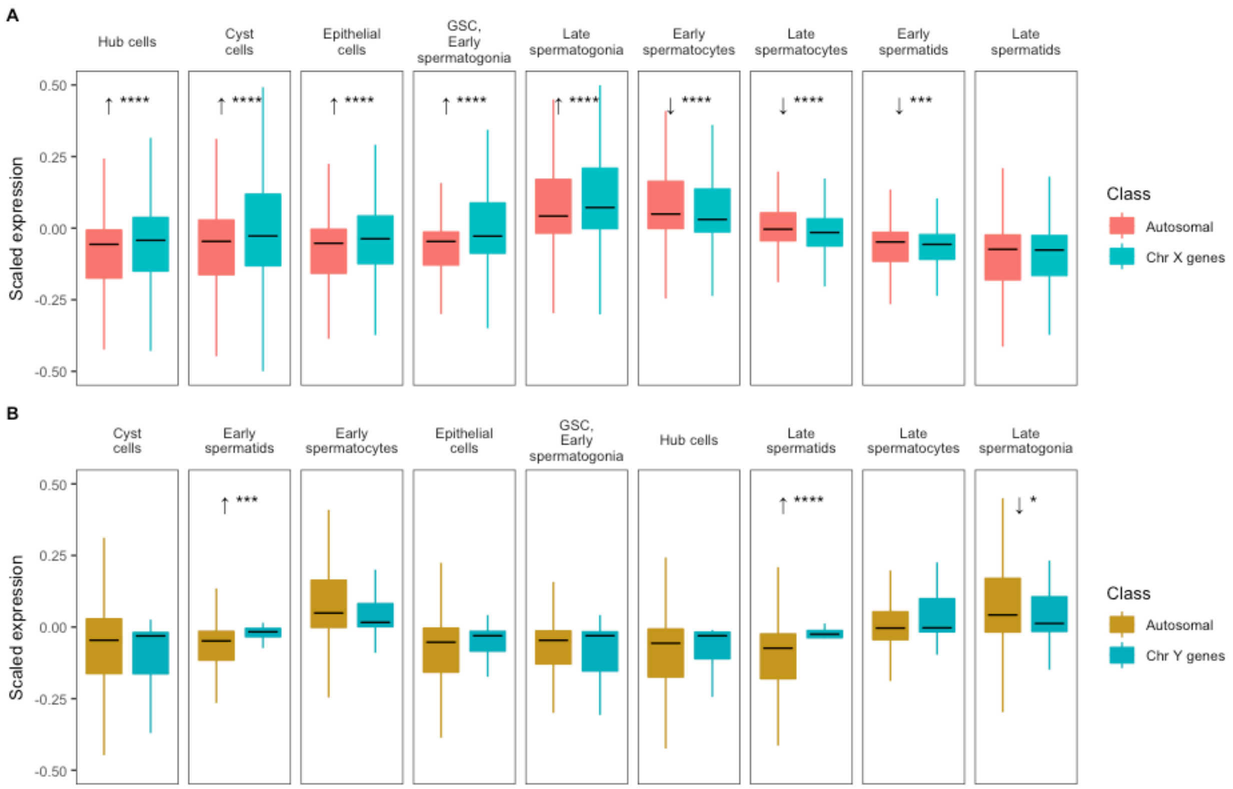


**Supplemental Figure 6: Scaled expression provides evidence of cell-type specific dosage compensation.** A.) Boxplots indicate the distribution of scaled expression of autosomal and X chromosome genes within each cell type. 0 represents a gene's mean expression across all cell types. In hub cells, epithelial cells, and premeiotic germ cells, scaled expression of X genes exceeds that of autosomal genes, suggesting that these cells experience X chromosome dosage compensation. Asterisks represent Holm-adjusted p values of directional Wilcoxon tests. B.) Scaled expression of Y chromosome genes exceeds that of the autosomes in late spermatocytes, early spermatids, and late spermatids. This indicates that after meiosis, Y chromosome genes are not downregulated to the same extent as autosomal genes. Asterisks represent p values as follows: ns: >0.05, *<0.05, **<0.005, ***<0.0005, ****<0.00005.

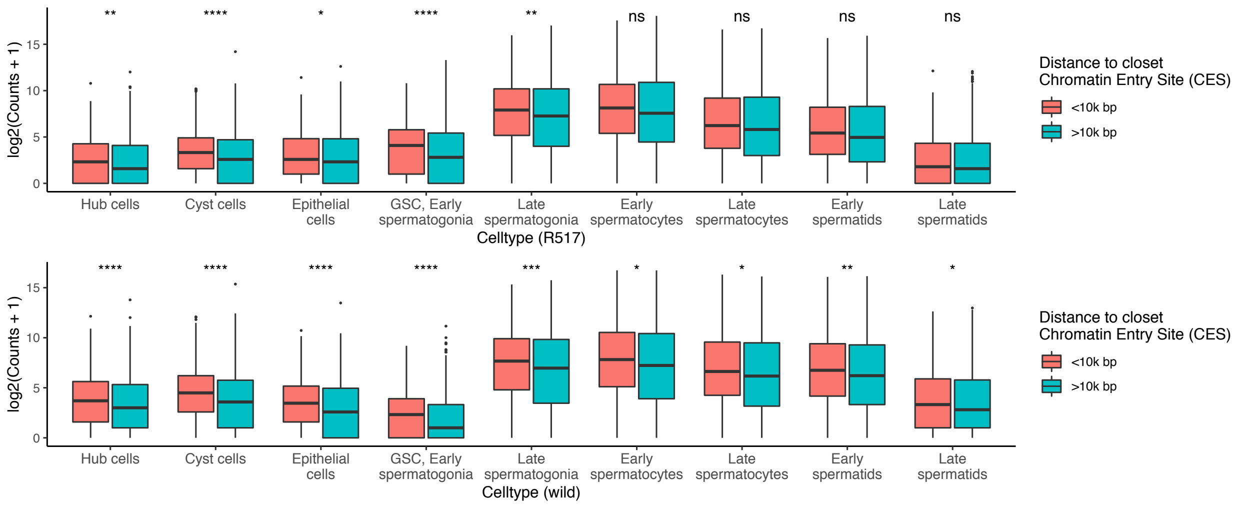


**Supplemental Figure 7: relationship between CES proximity and gene expression, grouped by cell type and strain.** Both datasets agree that somatic and pre-meiotic cells have a statistical enrichment of counts detected from genes within 10000 bp from a CES. In the wild dataset, however, proximal genes are enriched in every cell type, although less so in meiotic and post-meiotic cells than in pre-meiotic and somatic cells.

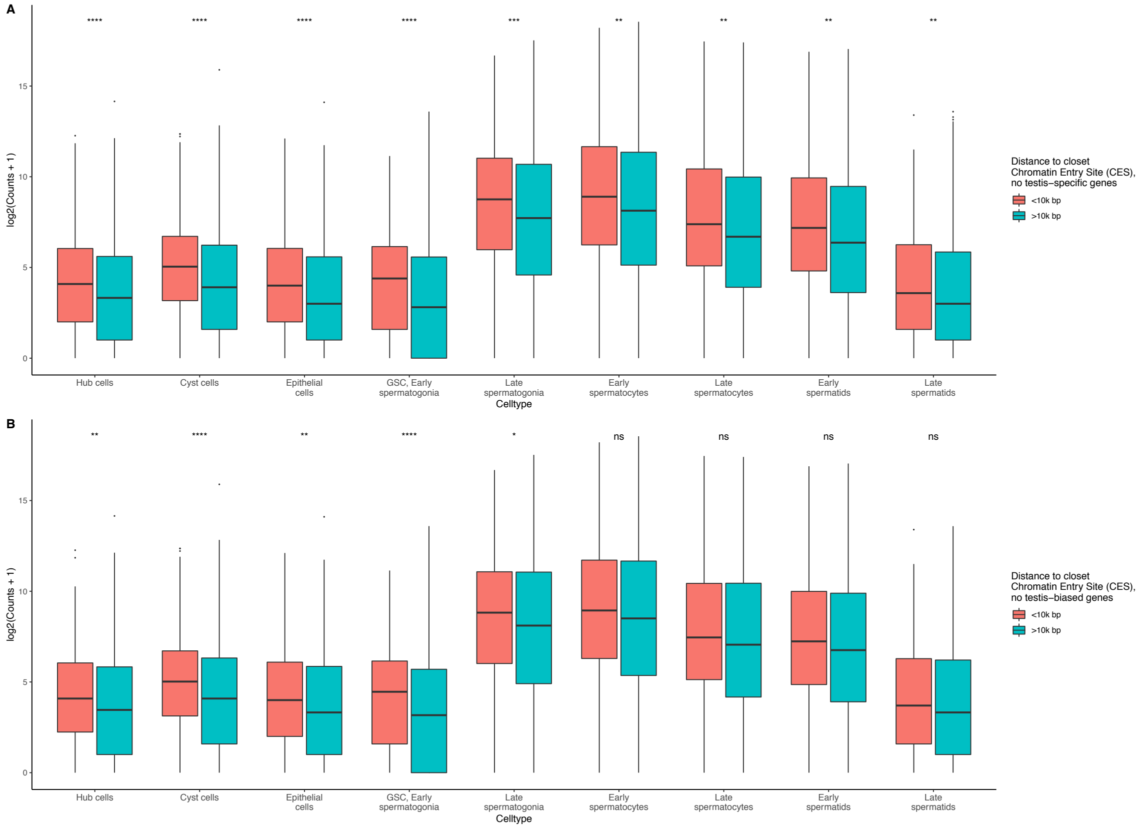


**Supplemental Figure 8: Figure 3 is not confounded by testis-specific or testis-biased genes**. Shown is the analysis from figure 4, repeated with testis-specific or testis-biased genes removed from the analysis. Neither result changes the conclusions in Figure 4.

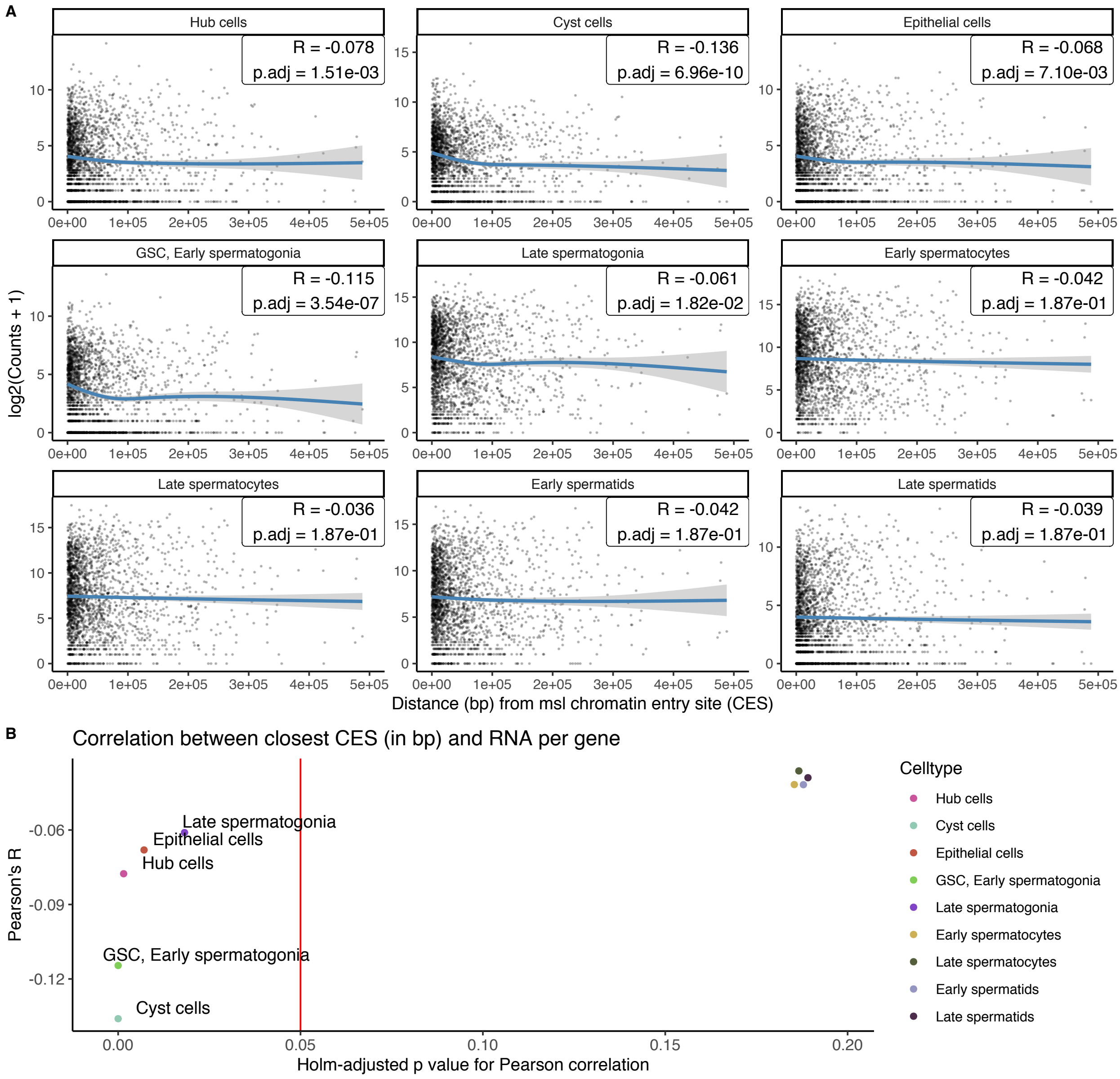


**Supplemental Figure 9: Close chromatin entry sites correlate with increased transcription of X chromosome genes in cell types experiencing DC.** A.) Each dot is an X chromosome gene, the X axis is the distance (in bp) between the gene start and the closest Chromatin Entry Site (CES) from Alekseyenko et al. 2008. For every gene, the Y axis is the log-transformed sum of all counts of that gene in a cell type. The black line is a Loess regression showing an approximate trend between the two axes. B.) Pearson's R shows that CES distance loosely correlates with RNA counts in hub, cyst, epithelial, GSC, early spermatogonia, and late spermatogonia cells, all cell types where we found evidence of dosage compensation. Spermatocytes and spermatids have Pearson's R closer to zero than DC-exhibiting cells, indicating less of a relationship between CES distance and transcription. In addition, these non-dosage-compensated cell types have a high Holm-adjusted p value, suggesting that distance and counts are not likely correlated in these cells.

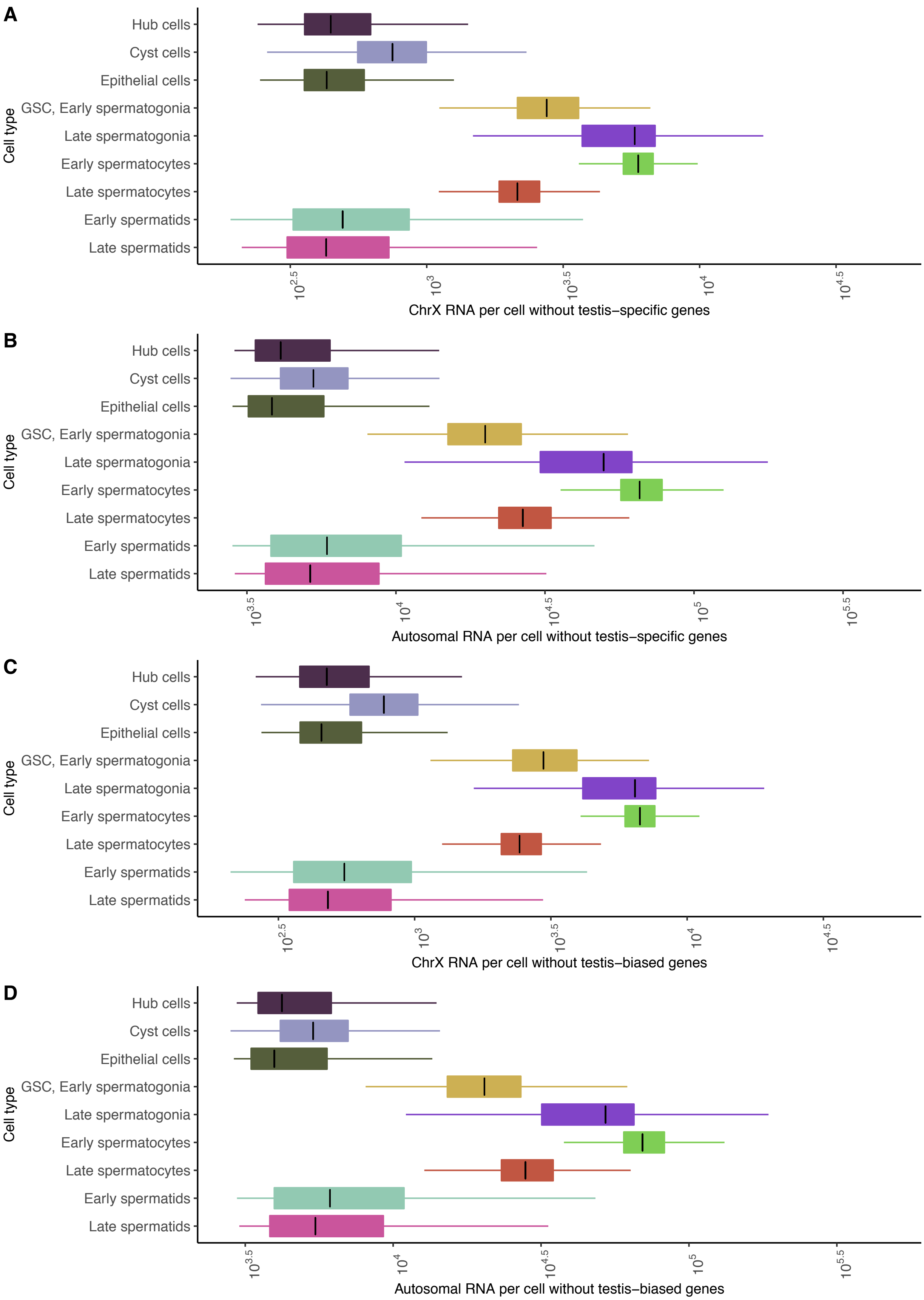
A
B
C
D
Cell type
Hub cells
Cyst cells
Epithelial cells
GSC, Early spermatogonia
Late spermatogonia
Early spermatocytes
Late spermatocytes
Early spermatids
Late spermatids
$10^{2.5}$
$10^{3}$
$10^{3.5}$
$10^{4}$
$10^{4.5}$
$10^{5}$
$10^{5.5}$
ChrX RNA per cell without testis–specific genes
Autosomal RNA per cell without testis–specific genes
ChrX RNA per cell without testis–biased genes
Autosomal RNA per cell without testis–biased genes

**Supplemental Figure 10: Figure 1 is not confounded by testis-specific or testis-biased genes**. Each panel corresponds to Figure 1B, with testis-specific or testis-biased genes removed from the dataset. Overall patterns of RNA per cell do not significantly change.

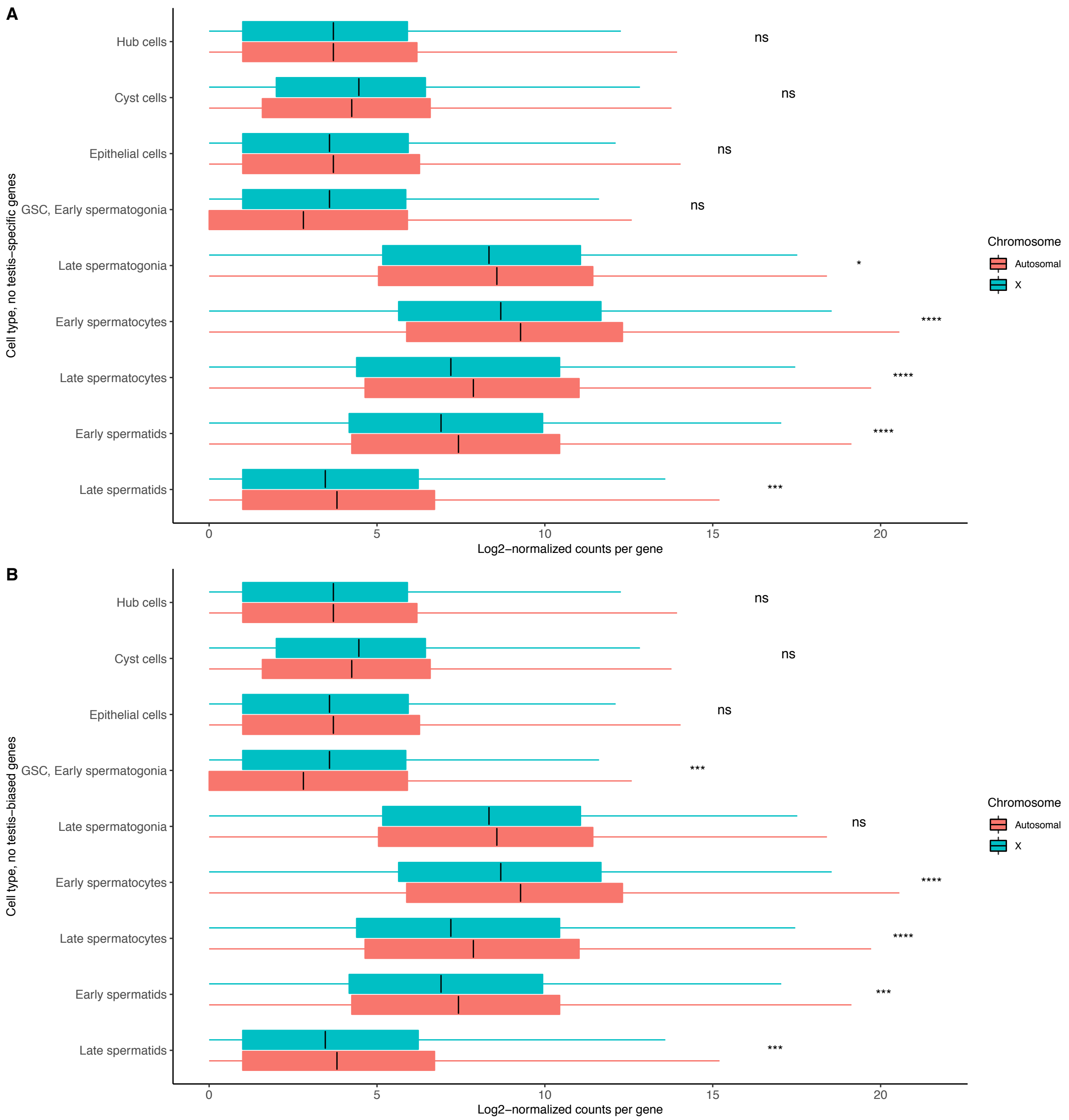


**Supplemental Figure 11: Testis-specific and testis-biased genes do not confound**

**the inference of pre-meiotic dosage compensation in figure 2.** This is the same analysis as Figure 2, with testis-specific genes removed. The overall patterns of pre-meiotic dosage compensation and meiotic X downregulation are preserved, indicating that these genes did not influence our main findings. One difference from figure 2 is that when testis-specific genes are removed, there is no longer statistical enrichment of X chromosome genes in GSC and early spermatogonia, indicating that testis-specific genes contribute to the appearance of X-chromosome overcompensation in these cells. Another difference is that without testis-biased genes, late spermatogonia no longer show statistical depletion of X chromosome counts, suggesting that these genes contribute to the earliest signs of reduced dosage compensation.

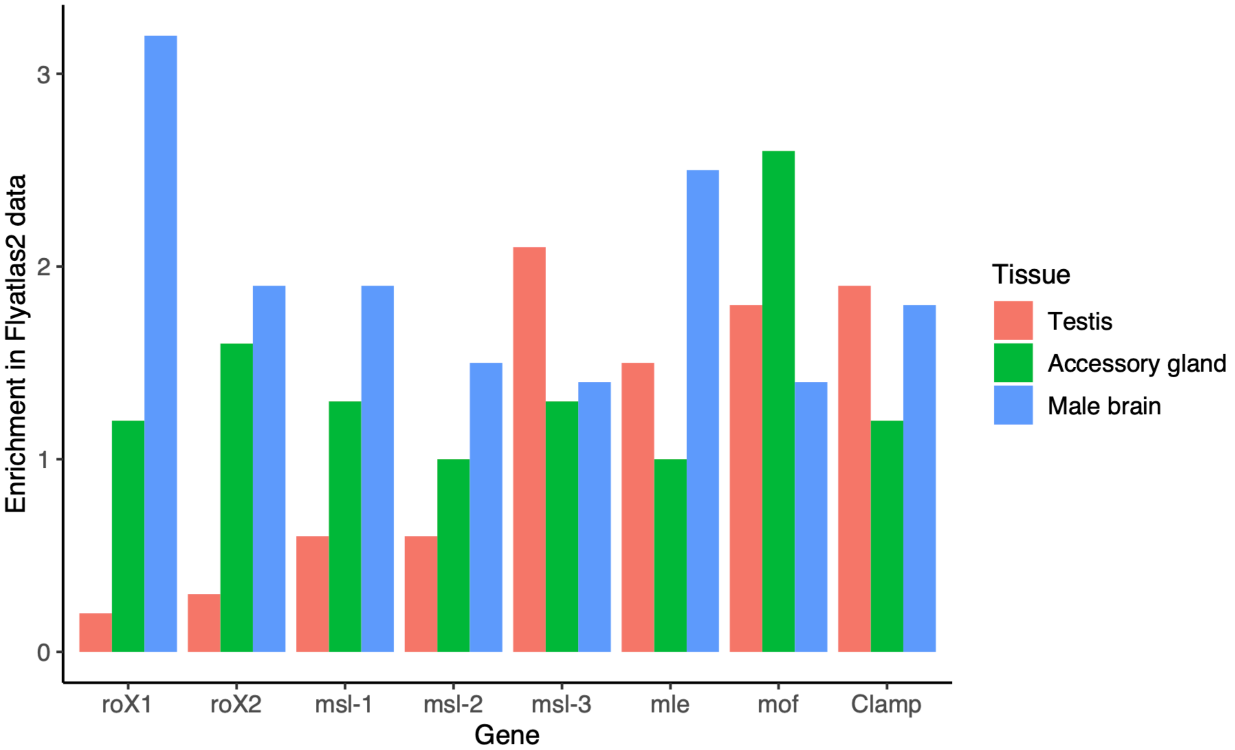


**Supplemental Figure 12: DCC gene enrichment in flyatlas2 data.** For each gene in the DCC, we queried the Flyatlas2 website for their calculated enrichments in selected tissues compared to all tissues. *RoX1*, *roX2, msl-1* and *msl-2* are very depleted in testis compared to somatic tissues, corresponding to their relatively stochastic expression in our sc-RNA-seq and FISH data.

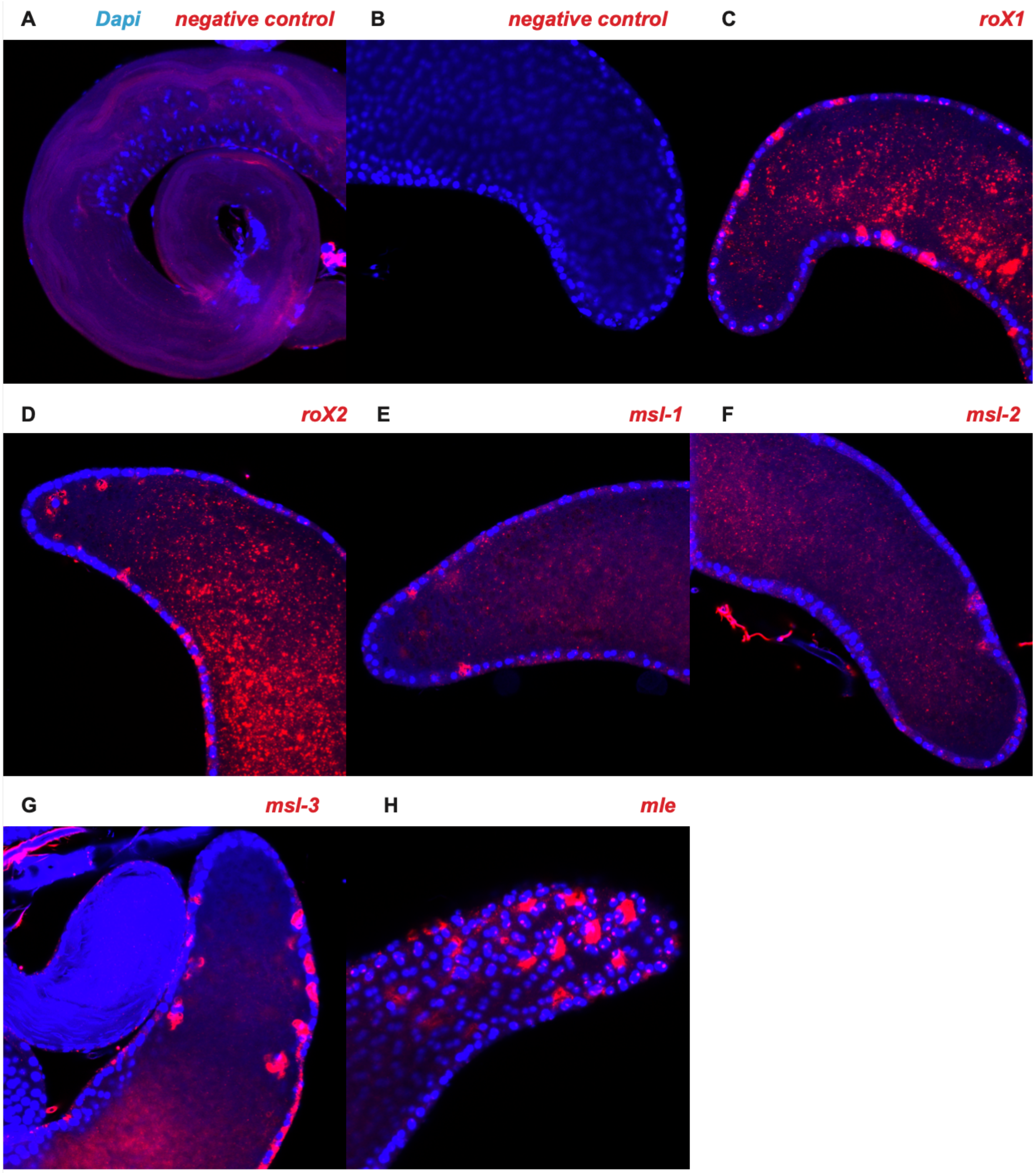


**Supplemental Figure 13: additional DCC gene FISH photos**. A) negative control in testis B) negative control in accessory gland C) *roX1* shows nuclear expression in accessory gland main cells and diffuse puncta in AG lumen, as well as membrane expression in secondary cells. D) In accessory gland, small patches of *roX2* are expressed in the cytoplasm and nuclei of MCs and along membranes of SCs. *roX2* is prevalent in AG lumen as diffuse foci. E) *mle* is expressed as dense foci localized in nuclei of MCs, with smaller foci in the cytoplasm of MCs. In SCs, it is enriched near membranes and shows smaller and fewer dots in the AG lumen. F) *msl-2* shows small foci distributed at low levels in MC cytoplasm. In SCs, it is enriched along membranes and in discrete foci in the AG lumen. G) *msl-3* shows strong expression patterns in the cytoplasm of MCs and the membranes of SCs, with smaller diffuse foci distributed in the AG lumen.

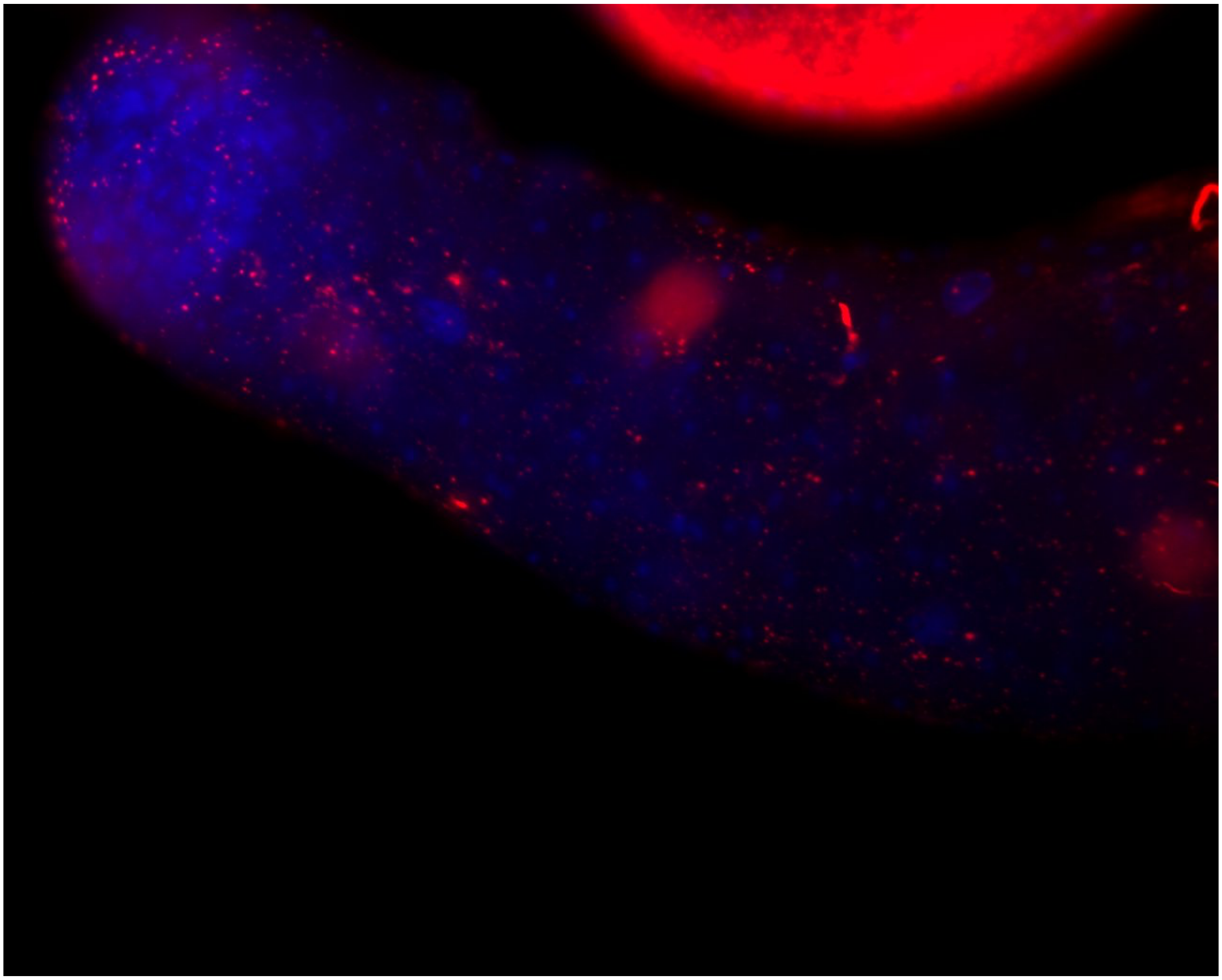

**Supplemental Figure 14: Enlarged RNA-FISH image of *msl-3*.** DAPI is shown in blue. In this image we see nuclear localization in hub and germline stem cells, nuclear and cytoplasmic expression in muscle and pigment cells, and cytoplasmic expression in cyst stem cells.

| Cell type | X counts | Autosome counts | X/autosome total count ratio |
|---|---|---|---|
| **Hub cells** | 662 | 6349 | 0.104 |
| **Cyst cells** | 895 | 6647 | 0.135 |
| **Epithelial** | 904 | 8669 | 0.104 |
| **GSC, early spermatogonia** | 3263 | 22448 | 0.145 |

| | | | |
|---|---|---|---|
| **Late spermatogonia** | 5979 | 49936 | 0.120 |
| **Early spermatocytes** | 6804 | 71689 | 0.095 |
| **Late spermatocytes** | 2749 | 31699 | 0.087 |
| **Early spermatids** | 785 | 8678 | 0.090 |
| **Late spermatids** | 851 | 9677 | 0.088 |

**Supplemental table 1:** Mean counts per cell from the X chromosome and autosomes in every cell type, corresponding to Figure 1B. The X chromosome is most highly transcribed compared to autosomes in GSC and spermatogonia.

| Cell type | P value | Pearson's R | Adjusted p value |
|---|---|---|---|
| **Hub cells*** | 2.15e-04 | 1.51e-03 | 1.51e-03 |
| **Cyst cells*** | 7.73e-11 | 6.96e-10 | 6.96e-10 |
| **Epithelial cells*** | 1.18e-03 | 7.10e-03 | 7.10e-03 |
| **GSC, early spermatogonia*** | 4.43e-08 | 3.54e-07 | 3.54e-07 |
| **Late spermatogonia*** | 3.64e-03 | 1.82e-02 | 1.82e-02 |
| **Early spermatocytes** | 4.71e-02 | 1.87e-01 | 1.87e-01 |
| **Late spermatocytes** | 8.48e-02 | 1.87e-01 | 1.87e-01 |
| **Early spermatids** | 4.68e-02 | 1.87e-01 | 1.87e-01 |
| **Late spermatids** | 6.37e-02 | 1.87e-01 | 1.87e-01 |

**Supplemental Table 2: Correlation between distance and transcription for X chromosome genes by cell type.** Corresponding to Supplemental figure 9, this is the Pearson's R between distance and log2(counts+1) for every gene per cell type, with Holm-adjusted p values. Cell types with evidence of dosage compensation are shown with an asterisk.

| Table | dm5 start | dm5 end | dm5 max | R5 coordinates | R6 coordinates |
|---|---|---|---|---|---|
| 1 | 366390 | 367190 | 366919 | X:366,390..367,190 | X:472,357..473,157 |
| 2 | 546603 | 549803 | 548347 | X:546,603..549,803 | X:652,570..655,770 |
| 3 | 655425 | 657601 | 656386 | X:655,425..657,601 | X:761,392..763,568 |
| 4 | 691829 | 692929 | 692277 | X:691,829..692,929 | X:797,796..798,896 |
| 5 | 936445 | 938345 | 937160 | X:936,445..938,345 | X:1,042,412..1,044,312 |
| 6 | 1267881 | 1269481 | 1268824 | X:1,267,881..1,269,481 | X:1,373,848..1,375,448 |

| | | | | | |
|---|---|---|---|---|---|
| 7 | 1356309 | 1357009 | 1356654 | X:1,356,309..1,357,009 | X:1,462,276..1,462,976 |
| 8 | 1374493 | 1375994 | 1375633 | X:1,374,493..1,375,994 | X:1,480,460..1,481,961 |
| 9 | 1587060 | 1588360 | 1588018 | X:1,587,060..1,588,360 | X:1,693,027..1,694,327 |
| 10 | 1774480 | 1775880 | 1775057 | X:1,774,480..1,775,880 | X:1,880,447..1,881,847 |
| 11 | 1786722 | 1788324 | 1787736 | X:1,786,722..1,788,324 | X:1,892,689..1,894,291 |
| 12 | 1918163 | 1919433 | 1918996 | X:1,918,163..1,919,433 | X:2,024,130..2,025,400 |
| 13 | 1955928 | 1957528 | 1956479 | X:1,955,928..1,957,528 | X:2,061,895..2,063,495 |
| 14 | 2072201 | 2074301 | 2072978 | X:2,072,201..2,074,301 | X:2,178,168..2,180,268 |
| 15 | 2125312 | 2126612 | 2125856 | X:2,125,312..2,126,612 | X:2,231,279..2,232,579 |
| 16 | 2236530 | 2238455 | 2237633 | X:2,236,530..2,238,455 | X:2,342,497..2,344,422 |
| 17 | 2339621 | 2341321 | 2340260 | X:2,339,621..2,341,321 | X:2,445,588..2,447,288 |
| 18 | 2492710 | 2493672 | 2493035 | X:2,492,710..2,493,672 | X:2,598,677..2,599,639 |
| 19 | 2520597 | 2521797 | 2521053 | X:2,520,597..2,521,797 | X:2,626,564..2,627,764 |
| 20 | 2628439 | 2629239 | 2628637 | X:2,628,439..2,629,239 | X:2,734,406..2,735,206 |
| 21 | 3375765 | 3377065 | NA | X:3,375,765..3,377,065 | X:3,481,732..3,483,032 |
| 22 | 3694068 | 3695885 | 3694893 | X:3,694,068..3,695,885 | X:3,800,035..3,801,852 |
| 23 | 3684985 | 3686792 | 3685816 | X:3,684,985..3,686,792 | X:3,790,952..3,792,759 |
| 24 | 3753268 | 3756637 | 3755452 | X:3,753,268..3,756,637 | X:3,859,235..3,862,604 |
| 25 | 3843793 | 3844593 | NA | X:3,843,793..3,844,593 | X:3,949,760..3,950,560 |
| 26 | 4020713 | 4022091 | 4021840 | X:4,020,713..4,022,091 | X:4,126,680..4,128,058 |
| 27 | 4438063 | 4439388 | 4439145 | X:4,438,063..4,439,388 | X:4,544,030..4,545,355 |
| 28 | 4578089 | 4578789 | 4578482 | X:4,578,089..4,578,789 | X:4,684,056..4,684,756 |
| 29 | 4598677 | 4600577 | 4599396 | X:4,598,677..4,600,577 | X:4,704,644..4,706,544 |
| 30 | 4969274 | 4970664 | 4970032 | X:4,969,274..4,970,664 | X:5,075,241..5,076,631 |
| 31 | 5308545 | 5310185 | 5309312 | X:5,308,545..5,310,185 | X:5,414,512..5,416,152 |
| 32 | 5563745 | 5565145 | 5564049 | X:5,563,745..5,565,145 | X:5,669,712..5,671,112 |
| 33 | 5653268 | 5654832 | 5653868 | X:5,653,268..5,654,832 | X:5,759,235..5,760,799 |
| 34 | 5681510 | 5684212 | 5682909 | X:5,681,510..5,684,212 | X:5,787,477..5,790,179 |
| 35 | 5804451 | 5806879 | 5806305 | X:5,804,451..5,806,879 | X:5,910,418..5,912,846 |
| 36 | 5976245 | 5977745 | 5977295 | X:5,976,245..5,977,745 | X:6,082,212..6,083,712 |
| 37 | 6122783 | 6123483 | NA | X:6,122,783..6,123,483 | X:6,228,750..6,229,450 |
| 38 | 6125383 | 6126429 | 6126019 | X:6,125,383..6,126,429 | X:6,231,350..6,232,396 |
| 39 | 6172809 | 6173997 | 6173582 | X:6,172,809..6,173,997 | X:6,278,776..6,279,964 |
| 40 | 6264168 | 6265668 | 6265067 | X:6,264,168..6,265,668 | X:6,370,135..6,371,635 |

| | | | | | |
|---|---|---|---|---|---|
| 41 | 6431763 | 6432463 | 6431910 | X:6,431,763..6,432,463 | X:6,537,730..6,538,430 |
| 42 | 6563384 | 6564901 | 6564053 | X:6,563,384..6,564,901 | X:6,669,351..6,670,868 |
| 43 | 6705400 | 6706845 | 6706224 | X:6,705,400..6,706,845 | X:6,811,367..6,812,812 |
| 44 | 7181094 | 7183678 | 7182556 | X:7,181,094..7,183,678 | X:7,287,061..7,289,645 |
| 45 | 7213876 | 7217359 | 7215106 | X:7,213,876..7,217,359 | X:7,319,843..7,323,326 |
| 46 | 7618472 | 7619472 | 7619172 | X:7,618,472..7,619,472 | X:7,724,439..7,725,439 |
| 47 | 7790451 | 7792008 | 7790905 | X:7,790,451..7,792,008 | X:7,896,418..7,897,975 |
| 48 | 7942825 | 7943937 | 7943529 | X:7,942,825..7,943,937 | X:8,048,792..8,049,904 |
| 49 | 7981841 | 7982605 | 7982237 | X:7,981,841..7,982,605 | X:8,087,808..8,088,572 |
| 50 | 8028174 | 8029974 | 8029113 | X:8,028,174..8,029,974 | X:8,134,141..8,135,941 |
| 51 | 8142104 | 8143004 | 8142580 | X:8,142,104..8,143,004 | X:8,248,071..8,248,971 |
| 52 | 8297145 | 8299963 | 8298407 | X:8,297,145..8,299,963 | X:8,403,112..8,405,930 |
| 53 | 8582587 | 8584087 | 8583332 | X:8,582,587..8,584,087 | X:8,688,554..8,690,054 |
| 54 | 8799008 | 8800137 | 8799773 | X:8,799,008..8,800,137 | X:8,904,975..8,906,104 |
| 55 | 9038744 | 9039544 | 9039023 | X:9,038,744..9,039,544 | X:9,144,711..9,145,511 |
| 56 | 9224784 | 9226114 | 9225778 | X:9,224,784..9,226,114 | X:9,330,751..9,332,081 |
| 57 | 9446480 | 9447780 | 9447253 | X:9,446,480..9,447,780 | X:9,552,447..9,553,747 |
| 58 | 9578393 | 9581795 | 9580787 | X:9,578,393..9,581,795 | X:9,684,360..9,687,762 |
| 59 | 9769835 | 9770835 | 9770616 | X:9,769,835..9,770,835 | X:9,875,802..9,876,802 |
| 60 | 9966323 | 9967023 | 9966601 | X:9,966,323..9,967,023 | X:10,072,290..10,072,990 |
| 61 | 10118520 | 10119764 | 10118946 | X:10,118,520..10,119,764 | X:10,224,487..10,225,731 |
| 62 | 10276207 | 10277160 | 10276600 | X:10,276,207..10,277,160 | X:10,382,174..10,383,127 |
| 63 | 10357313 | 10358390 | 10357608 | X:10,357,313..10,358,390 | X:10,463,280..10,464,357 |
| 64 | 10367323 | 10368423 | 10368421 | X:10,367,323..10,368,423 | X:10,473,290..10,474,390 |
| 65 | 10665610 | 10666474 | 10665977 | X:10,665,610..10,666,474 | X:10,771,577..10,772,441 |
| 66 | 10745605 | 10746405 | 10745748 | X:10,745,605..10,746,405 | X:10,851,572..10,852,372 |
| 67 | 10758754 | 10760630 | 10759843 | X:10,758,754..10,760,630 | X:10,864,721..10,866,597 |
| 68 | 10815961 | 10816662 | NA | X:10,815,961..10,816,662 | X:10,921,928..10,922,629 |
| 69 | 11038326 | 11039763 | 11039116 | X:11,038,326..11,039,763 | X:11,144,293..11,145,730 |
| 70 | 11057400 | 11058524 | 11057415 | X:11,057,400..11,058,524 | X:11,163,367..11,164,491 |
| 71 | 11290632 | 11293111 | 11291584 | X:11,290,632..11,293,111 | X:11,396,599..11,399,078 |
| 72 | 11293774 | 11294576 | NA | X:11,293,774..11,294,576 | X:11,399,741..11,400,543 |
| 73 | 11472953 | 11474927 | 11474270 | X:11,472,953..11,474,927 | X:11,578,920..11,580,894 |
| 74 | 11595401 | 11598135 | 11595759 | X:11,595,401..11,598,135 | X:11,701,368..11,704,102 |

| | | | | | |
|---|---|---|---|---|---|
| 75 | 11616668 | 11617791 | 11617352 | X:11,616,668..11,617,791 | X:11,722,635..11,723,758 |
| 76 | 11722160 | 11723229 | 11722824 | X:11,722,160..11,723,229 | X:11,828,127..11,829,196 |
| 77 | 11758511 | 11761239 | 11760360 | X:11,758,511..11,761,239 | X:11,864,478..11,867,206 |
| 78 | 11904033 | 11906110 | 11905008 | X:11,904,033..11,906,110 | X:12,010,000..12,012,077 |
| 79 | 12542747 | 12545401 | 12544973 | X:12,542,747..12,545,401 | X:12,648,714..12,651,368 |
| 80 | 12604385 | 12605085 | 12604739 | X:12,604,385..12,605,085 | X:12,710,352..12,711,052 |
| 81 | 12609483 | 12610183 | 12610103 | X:12,609,483..12,610,183 | X:12,715,450..12,716,150 |
| 82 | 12646261 | 12647087 | NA | X:12,646,261..12,647,087 | X:12,752,228..12,753,054 |
| 83 | 12653827 | 12656232 | 12654565 | X:12,653,827..12,656,232 | X:12,759,794..12,762,199 |
| 84 | 12804471 | 12806996 | 12805027 | X:12,804,471..12,806,996 | X:12,910,438..12,912,963 |
| 85 | 13093353 | 13096385 | 13095036 | X:13,093,353..13,096,385 | X:13,199,320..13,202,352 |
| 86 | 13157101 | 13157829 | 13157555 | X:13,157,101..13,157,829 | X:13,263,068..13,263,796 |
| 87 | 13234896 | 13237464 | 13235886 | X:13,234,896..13,237,464 | X:13,340,863..13,343,431 |
| 88 | 13283549 | 13284273 | NA | X:13,283,549..13,284,273 | X:13,389,516..13,390,240 |
| 89 | 13314325 | 13315325 | 13315226 | X:13,314,325..13,315,325 | X:13,420,292..13,421,292 |
| 90 | 13630255 | 13631714 | 13630460 | X:13,630,255..13,631,714 | X:13,736,222..13,737,681 |
| 91 | 13664247 | 13665281 | 13664853 | X:13,664,247..13,665,281 | X:13,770,214..13,771,248 |
| 92 | 13720782 | 13721482 | 13721027 | X:13,720,782..13,721,482 | X:13,826,749..13,827,449 |
| 93 | 13889981 | 13891745 | 13890982 | X:13,889,981..13,891,745 | X:13,995,948..13,997,712 |
| 94 | 13996764 | 14008829 | 13998406 | X:13,996,764..14,008,829 | X:14,102,731..14,114,796 |
| 95 | 14010329 | 14012229 | 14011574 | X:14,010,329..14,012,229 | X:14,116,296..14,118,196 |
| 96 | 14720985 | 14723179 | 14722568 | X:14,720,985..14,723,179 | X:14,826,952..14,829,146 |
| 97 | 14944690 | 14947962 | 14946204 | X:14,944,690..14,947,962 | X:15,050,657..15,053,929 |
| 98 | 14979863 | 14981953 | 14980466 | X:14,979,863..14,981,953 | X:15,085,830..15,087,920 |
| 99 | 15477142 | 15479342 | 15478359 | X:15,477,142..15,479,342 | X:15,583,109..15,585,309 |
| 100 | 15623738 | 15625770 | 15624768 | X:15,623,738..15,625,770 | X:15,729,705..15,731,737 |
| 101 | 15693722 | 15696722 | 15695055 | X:15,693,722..15,696,722 | X:15,799,689..15,802,689 |
| 102 | 15728080 | 15729132 | 15728442 | X:15,728,080..15,729,132 | X:15,834,047..15,835,099 |
| 103 | 15754121 | 15755823 | 15754707 | X:15,754,121..15,755,823 | X:15,860,088..15,861,790 |
| 104 | 15769177 | 15770977 | 15769834 | X:15,769,177..15,770,977 | X:15,875,144..15,876,944 |
| 105 | 15889446 | 15891074 | 15890650 | X:15,889,446..15,891,074 | X:15,995,413..15,997,041 |
| 106 | 16167399 | 16168907 | 16168205 | X:16,167,399..16,168,907 | X:16,273,366..16,274,874 |
| 107 | 16203823 | 16204705 | 16204038 | X:16,203,823..16,204,705 | X:16,309,790..16,310,672 |
| 108 | 16250181 | 16251481 | 16250882 | X:16,250,181..16,251,481 | X:16,356,148..16,357,448 |

| | | | | | |
|---|---|---|---|---|---|
| 109 | 16455838 | 16456864 | NA | X:16,455,838..16,456,864 | X:16,561,805..16,562,831 |
| 110 | 16501012 | 16502877 | 16502037 | X:16,501,012..16,502,877 | X:16,606,979..16,608,844 |
| 111 | 16621477 | 16623077 | NA | X:16,621,477..16,623,077 | X:16,727,444..16,729,044 |
| 112 | 16689604 | 16692767 | 16692394 | X:16,689,604..16,692,767 | X:16,795,571..16,798,734 |
| 113 | 16775983 | 16776983 | 16776671 | X:16,775,983..16,776,983 | X:16,881,950..16,882,950 |
| 114 | 16969751 | 16971404 | 16970798 | X:16,969,751..16,971,404 | X:17,075,718..17,077,371 |
| 115 | 17030560 | 17033492 | 17031424 | X:17,030,560..17,033,492 | X:17,136,527..17,139,459 |
| 116 | 17048415 | 17049615 | 17048811 | X:17,048,415..17,049,615 | X:17,154,382..17,155,582 |
| 117 | 17178939 | 17180238 | 17179865 | X:17,178,939..17,180,238 | X:17,284,906..17,286,205 |
| 118 | 17538938 | 17540038 | NA | X:17,538,938..17,540,038 | X:17,644,905..17,646,005 |
| 119 | 17547893 | 17550393 | 17549607 | X:17,547,893..17,550,393 | X:17,653,860..17,656,360 |
| 120 | 17598986 | 17601152 | 17600087 | X:17,598,986..17,601,152 | X:17,704,953..17,707,119 |
| 121 | 17714956 | 17717075 | 17715734 | X:17,714,956..17,717,075 | X:17,820,923..17,823,042 |
| 122 | 17819963 | 17821239 | NA | X:17,819,963..17,821,239 | X:17,925,930..17,927,206 |
| 123 | 17987153 | 17988753 | 17987981 | X:17,987,153..17,988,753 | X:18,093,120..18,094,720 |
| 124 | 17991993 | 17992693 | 17992605 | X:17,991,993..17,992,693 | X:18,097,960..18,098,660 |
| 125 | 18268228 | 18269396 | 18268774 | X:18,268,228..18,269,396 | X:18,374,195..18,375,363 |
| 126 | 18388748 | 18390439 | 18389692 | X:18,388,748..18,390,439 | X:18,494,715..18,496,406 |
| 127 | 18547720 | 18549220 | 18548520 | X:18,547,720..18,549,220 | X:18,653,687..18,655,187 |
| 128 | 18683463 | 18685163 | 18684116 | X:18,683,463..18,685,163 | X:18,789,430..18,791,130 |
| 129 | 18742623 | 18744205 | 18743223 | X:18,742,623..18,744,205 | X:18,848,590..18,850,172 |
| 130 | 18781253 | 18782810 | 18782227 | X:18,781,253..18,782,810 | X:18,887,220..18,888,777 |
| 131 | 19089662 | 19091279 | 19090428 | X:19,089,662..19,091,279 | X:19,195,629..19,197,246 |
| 132 | 19166379 | 19167579 | 19167145 | X:19,166,379..19,167,579 | X:19,272,346..19,273,546 |
| 133 | 19383500 | 19384800 | 19384312 | X:19,383,500..19,384,800 | X:19,489,467..19,490,767 |
| 134 | 19471825 | 19473867 | 19473298 | X:19,471,825..19,473,867 | X:19,577,792..19,579,834 |
| 135 | 19518350 | 19519250 | 19518692 | X:19,518,350..19,519,250 | X:19,624,317..19,625,217 |
| 136 | 19534337 | 19535866 | 19535151 | X:19,534,337..19,535,866 | X:19,640,304..19,641,833 |
| 137 | 19582304 | 19584399 | 19583395 | X:19,582,304..19,584,399 | X:19,688,271..19,690,366 |
| 138 | 19624437 | 19625391 | NA | X:19,624,437..19,625,391 | X:19,730,404..19,731,358 |
| 139 | 19635080 | 19635856 | 19635374 | X:19,635,080..19,635,856 | X:19,741,047..19,741,823 |
| 140 | 19752914 | 19753814 | 19753490 | X:19,752,914..19,753,814 | X:19,858,881..19,859,781 |
| 141 | 19917716 | 19919805 | 19918722 | X:19,917,716..19,919,805 | X:20,023,683..20,025,772 |
| 142 | 20052603 | 20053719 | 20053190 | X:20,052,603..20,053,719 | X:20,158,570..20,159,686 |

| | | | | | |
|---|---|---|---|---|---|
| 143 | 20102872 | 20103676 | NA | X:20,102,872..20,103,676 | X:20,231,798..20,232,602 |
| 144 | 20282194 | 20283326 | 20282867 | X:20,282,194..20,283,326 | X:20,411,167..20,412,299 |
| 145 | 20920307 | 20923420 | 20921199 | X:20,920,307..20,923,420 | X:21,049,280..21,052,393 |
| 146 | 21102348 | 21103548 | 21103032 | X:21,102,348..21,103,548 | X:21,231,321..21,232,521 |
| 147 | 21187950 | 21189650 | 21188853 | X:21,187,950..21,189,650 | X:21,316,923..21,318,623 |
| 148 | 21239454 | 21241240 | 21240544 | X:21,239,454..21,241,240 | X:21,368,427..21,370,213 |
| 149 | 21869967 | 21871175 | 21870626 | X:21,869,967..21,871,175 | X:22,468,131..22,469,339 |
| 150 | 21936497 | 21938643 | 21937487 | X:21,936,497..21,938,643 | X:22,534,661..22,536,807 |

**Supplementary Table 3: Chromatin entry sites from Alekseyenko et. al 2008 (DOI: 10.1016/j.cell.2008.06.033)**. We have converted the start and end coordinates to FlyBase R6 version using the FlyBase coordinates converter.
